# TGF-β2 regulates neuronal Ankyrin-G and promotes its interaction with KCC2

**DOI:** 10.1101/2023.12.05.570078

**Authors:** Anastasia Rigkou, Abhishek Pethe, Eleni Roussa

**Affiliations:** Institute for Anatomy and Cell Biology, Department of Molecular Embryology, Faculty of Medicine, Albert-Ludwigs-Universität Freiburg, Albertstr. 17, D-79104 Freiburg, Germany

**Keywords:** growth factors, axon initial segment, potassium-chloride-cotransporter, scaffolding protein, brainstem

## Abstract

The neuronal K^+^/Cl^-^ cotransporter 2 (KCC2) is the major Cl^-^ extruder in CNS neurons and responsible for fast hyperpolarizing postsynaptic inhibition in mature neurons. Impaired KCC2 function has been associated with several brain pathologies. KCC2 forms immunocomplexes with several proteins that may regulate KCC2 membrane trafficking, stability and function, thus, tuning important cellular processes, including chloride homeostasis and dendritic spine development. In the brain, the scaffold protein Ankyrin-G, encoded by the *Ank3* gene, is expressed in several isoforms with distinct spatial and temporal expression patterns, is regulated by TGF-β signalling and is proposed as a KCC2 interaction partner. Moreover, *Ank3* gene has been implicated in several neuropsychiatric disorders.

Here, we investigated a putative impact of transforming growth factor beta 2 (TGF-β2) on KCC2/Ankyrin-G interaction using quantitative RT-PCR, immunoblotting, immunoprecipitation and immunofluorescence in mouse immature and differentiated hippocampal neurons and in forebrain and brainstem tissue from *Tgf-β2* deficient mice. The results show TGF-β2-dependent downregulation of *Ank3* transcripts, as well as KCC2/Ankyrin-G interaction in mouse brainstem tissue at embryonic day (E) 17.5. *In vitro*, loss of *Tgf-β2* resulted in significantly reduced axonal and somatic Ankyrin-G in immature neurons and significantly reduced somatic Ankyrin-G abundance in differentiated mouse hippocampal neurons. Membrane abundance of Ankyrin-G was downregulated in *Tgf-β2* mutants as well, a phenotype rescued by application of exogenous TGF-β2. Moreover, the results suggest the presence of a Golgi-associated Ankyrin-G isoform in neurons and significantly impaired membrane KCC2 abundance following knock down of *Ank3*. Thus, the present study provides new insights into Ankyrin-G regulation by TGF-β2 in neurons and first evidence of a TGF-β2-regulated interaction of KCC2 with Ankyrin-G. Moreover, these results strengthen the notion for TGF-β2 as pivotal regulator of KCC2 abundance and function.

## INTRODUCTION

Mature CNS neurons primarily rely on K^+^/Cl^-^ cotransporter 2 (KCC2; encoded by *Slc12a5* gene) as the main mechanism for chloride extrusion. Over the course of neuronal development, the upregulation of KCC2 expression and activity plays a crucial role in the transition of GABAergic responses from excitatory to inhibitory (Rivera et al., 1999). Genetic mutations of *Kcc2* in humans have been associated with the development of seizures and epilepsy (Puskarjov et al., 2014; Kahle et al., 2014; Stödberg et al., 2015; Saitsu et al., 2016), while *Kcc2* variants were found in patients with autism and schizophrenia (Merner et al., 2015). Additionally, several studies have suggested that impaired expression and function of KCC2 are implicated in the pathogenesis of various neurological disorders such as neuropathic pain (Mapplebeck et al., 2019), ischemia (Jaenisch et al., 2010) and autism (Tyzio et al., 2014).

Besides its involvement in neuronal chloride homeostasis, KCC2 has been associated with non-canonical, i.e. Cl^-^ transport-independent functions, such as dendritic spine morphogenesis, maintenance and plasticity at excitatory synapses (Li et al., 2007; Fiumelli et al., 2013; Llano et al., 2015; Mavrovic et al., 2020). In neurons lacking *Kcc2*, the number of functional synapses was decreased, while dendritic spine morphology appeared mainly immature and filopodia-like. This phenotype was restored by the expression of KCC2 lacking the N-terminal domain, responsible for transport function (Li et al., 2007).

KCC2 forms complexes with numerous types of proteins that may regulate its membrane trafficking, stability and function, thus, tuning important cellular processes, including chloride homeostasis and dendritic spine development. Interaction of KCC2 with Rab11b downstream of TGF-β2/CREB signaling regulates its membrane trafficking and functionality (Roussa et al., 2016). Furthermore, it was reported that KCC2 interacts with the scaffolding protein Gephyrin, which promotes its membrane expression and clustering near GABAergic synapses, thereby regulating its transport activity (Awabdh et al., 2022). The non-canonical function of KCC2 is mediated through its interaction with the spine cytoskeleton, particularly the cytoskeleton-associated protein 4.1N (Li et al., 2007) and the guanine nucleotide exchange factor β-PIX (Llano et al., 2015).

Recently, systematic KCC2 interactome studies have been performed and unveiled several putative KCC2 interaction partners (Smalley et al., 2020a; 2020b; Mahadevan et al., 2017). Ankyrin-B and Ankyrin-G were proposed as KCC2 interacting proteins in plasma membranes from adult mouse forebrain (Smalley et al., 2020b). Ankyrins are scaffolding proteins that link plasma membrane proteins to the actin/β-spectrin cytoskeleton. *Ank3* gene, encoding Ankyrin-G, has been implicated in a spectrum of neuropsychiatric disorders such as bipolar disorder and autism (Ferreira et al., 2008; Schulze et al., 2009; Bi et al., 2012). Human *Ank3* gene codes for different isoforms of Ankyrin-G with distinct spatial and temporal expression patterns and functions in the brain (reviewed by Yoon et al., 2022). Three of these isoforms are collectively known as Ankyrin-G 190 kDa because they migrate at ∼190 kDa when analysed by SDS-PAGE. Structurally, Ankyrin-G 190 kDa isoform(s) consist(s) of an ankyrin-repeat domain, a spectrin-binding domain and a regulatory domain. The 190 kDa isoform of Ankyrin-G is enriched in dendrites, regulating dendritic spine morphology (Smith et al., 2014). A fourth isoform contains an additional large exon. Alternative splicing of this large exon gives rise to two variants known as Ankyrin-G 270/480 kDa giant isoforms which are typically found in the axon initial segment and nodes of Ranvier where they play a critical role in organizing proteins, including ion channels, transporters and cell adhesion molecules (Jenkins et al., 2015). Smaller Ankyrin-G isoforms with molecular weight of 110 kDa, lack the ankyrin-repeat domain and have been associated with the regulation of endocytosis and lysosomal degradation (Ignatiuk et al., 2006).

Transforming growth factor betas (TGF-βs) are members of the TGF-β superfamily of signaling molecules, with pleiotropic functions in different cell types regulating development and differentiation. In the CNS, TGF-βs regulate neuronal survival and apoptosis during development, and have been shown to have effects on synaptogenesis, neural network function and neuronal plasticity (reviewed by Krieglstein et al., 2011; Meyers and Kessler, 2017). TGF-βs consist of three isoforms (TGF-β1,-β2,-β3) with distinct expression patterns in the nervous system (Unsicker et al., 1991). While constitutive loss of individual TGF-β genes are lethal in mice, the observed phenotypes are unique and isoform-specific (Dünker and Krieglstein, 2000). *Tgf-β2* constitutive knockout mice die at birth due to congenital cyanosis (Sanford et al., 1997) and exhibit abnormal excitatory and inhibitory postsynaptic currents in pre-Bötzinger complex neurons in the brainstem, demonstrating the significant role of the TGF-β2 isoform in neuronal development and synaptic transmission (Heupel et al., 2008). We have previously reported multiple modes of regulation of KCC2 by TGF-β2 throughout neuronal development. In immature neurons TGF-β2 regulates *Kcc2* transcription via transcription factor AP2β (Rigkou et al., 2022), whereas in differentiating and mature neurons it regulates the (de)phosphorylation of KCC2 at T1007 and membrane trafficking and functionality of KCC2 through a CREB/Rab11b mechanism (Roussa et al., 2016; Rigkou et al., 2022). Another study demonstrated that TGF-β signaling promotes dendritic spine development in cortical neurons by stabilizing Ankyrin-G through interaction with the deubiquitinase Usp9X (Yoon et al., 2020).

Taken into consideration the context-dependent TGF-β2 mode of action, and based on the background above, in the present study, we sought to investigate putative TGF-β2-dependent effects on KCC2/AnkG interaction. The results show TGF-β2-dependent regulation of *Ank3* transcripts, as well as KCC2/Ankyrin-G interaction in mouse brainstem tissue at embryonic day E17.5. *In vitro*, loss of *Tgf-β2* resulted in reduced somatic and axonal Ankyrin-G in immature neurons and reduced somatic and membrane Ankyrin-G abundance in differentiated mouse hippocampal neurons. Moreover, the results support a Golgi-associated Ankyrin-G isoform in neurons and significantly impaired membrane KCC2 abundance following knock down of *Ank3*. In summary, the present study provides new insights into Ankyrin-G regulation by TGF-β2 and first evidence of a direct or indirect TGF-β2-regulated interaction of KCC2 with Ankyrin-G.

## MATERIALS & METHODS

### Antibodies and reagents/chemicals

The primary antibodies used in co-immunoprecipitation, immunoblotting and immunocytochemistry are the following: rabbit polyclonal anti-KCC2 (Millipore Cat# 07-432, RRID:AB_310611), rabbit polyclonal anti-ankyrin-G (Synaptic Systems Cat# 386 003, RRID:AB_2661876), mouse monoclonal anti-ankyrin-G (Santa Cruz Biotechnology Cat# sc-12719, RRID:AB_626674), rabbit polyclonal anti-GOLGA5 (GeneTex Cat# GTX104255, RRID:AB_2037117) and mouse monoclonal anti-GAPDH (Proteintech Cat# 60004-1-Ig, RRID:AB_2107436). For immunoblotting, horse anti-mouse (Cell Signaling Technology Cat# 7076, RRID:AB_330924) or anti-rabbit (Cell Signaling Technology Cat# 7074, RRID:AB_2099233) IgG coupled to horseradish peroxidase were used as secondary antibodies. For immunofluorescence, donkey anti-rabbit AlexaFluor594 (Jackson ImmunoResearch Labs Cat# 711-585-152, RRID:AB_2340621) and donkey anti-mouse AlexaFluor488 (Jackson ImmunoResearch Labs Cat# 715-545-150, RRID:AB_2340846) were used as secondary antibodies. Recombinant human TGF-β2 (Cat# 302-B2-002/CF) was purchased from R&D Systems.

### Animals

All protocols were carried out in accordance with German ethical guidelines for laboratory animals and approved by the Institutional Animal Care and Use Committee of the University of Freiburg and the ethics committee of the City of Freiburg (authorizations: X19/09C, X22/11B and G-21/140). Adult C57BL/6N mice (strain code 027) of either sex were maintained on a 12 h dark/light cycle with food and water ad libitum. Mice were sacrificed by cervical dislocation and all efforts were made to minimize suffering. Embryos were sacrificed by decapitation at embryonic day (E) 17.5 of gestation, brains were isolated and processed as described below. Heterozygous *Tgf-β2*^+/-^ mice were crossed to obtain *Tgf-β2* deficient mice (*Tgf-β2*^-/-^) which have been described earlier (Sanford et al., 1997).

### Culture of mouse primary hippocampal neurons

Hippocampi were isolated from C57BL/6N, *Tgf-β2^+/+^* and *Tgf-β2^-/-^* mice at E17.5 and incubated in 0.25% trypsin for 20 min at 37 °C, followed by gentle trituration using silicon coated pasteur pipettes, as previously described (Lacmann et al., 2007; Roussa et al., 2016). Dissociated cells were resuspended in NeuroBasal medium supplemented with 2% B27, 1% PSN, 5 µg/ml Apo-transferin, 0.5 M L-glutamin, 0.8 µg/ml superoxide dismutase, 1 µg/ml glutathion and plated either onto poly-L-ornithin/ laminin coated 12-mm^2^ glass coverslips in 24-well plates at a density of 50.000-80.000 cells/coverslip or on 6-well-plates at a density of 10^6^ cells/well. At day in vitro (DIV) 4 and 8 half the medium was replaced with fresh medium containing 1 µg/ml and 0.5 µg/ml AraC (Cytosine β-D-arabinofuranoside) respectively, to reduce proliferation of glial cells. The neurons were treated at DIV4 or 12 with 2 ng/mL recombinant human TGF-β2 for 60 minutes and processed for immunoblotting or immunocytochemistry.

### Transfection of mouse primary hippocampal neurons

Mouse primary hippocampal neurons grown on coverslips were transiently transfected at DIV11 with Alexa 488-labeled siRNA either specifically targeting mouse *Kcc2b* (Mm_Slc12a5_1 FlexiTube siRNA, GeneGlobe ID: SI01419243, GenBank accession no: NM_020333; Qiagen) or *Ank3* mRNA (Mm_Ank3_3 FlexiTube siRNA, GeneGlobe ID: SI00898009, GenBank accession no: NM_009670, NM_146005, NM_170687, NM_170688, NM_170689, NM_170690, NM_170728, NM_170729, NM_170730; Qiagen) or with control negative siRNA, a sequence that reveals no homology with any known mammalian gene (AllStars Negative Control siRNA, cat. no. 1027292, Qiagen), as previously described (Oehlke et al., 2011). Per coverslip, 2 μL of HiPerfect transfection reagent (cat. no. 301705, Qiagen) were mixed with 100 μL NeuroBasal medium without supplements. Subsequently, 100 ng siRNA was added followed by incubation in the dark for 30 min to allow the formation of the complexes, before being pipetted onto the cells. 24h post transfection the cells were fixed and processed for immunocytochemistry.

### Crude membrane preparation

Crude membranes from adult mouse brain tissue were obtained as described previously (Pethe et al., 2023). Briefly, adult mice brains were isolated and homogenized in an isolation buffer (250mM sucrose, 40 mM MOPS, 0.1 mM EGTA, 0.1 mM MgSO4, pH 7.4) using a homogenizer (IKA-RW15) at 11,000 RPM with at least 20 up-and-down motions, on ice. The homogenate was centrifuged at 1,500 x g for 10 min to remove unlysed debris. The supernatant was collected and centrifuged at 1,000 x g for 10 min. The pellet was discarded and further centrifuged at 4,000 x g for 10 min. The supernatant thus obtained was centrifuged in an ultracentrifuge (Beckmann-Coulter XPN90, SW-41Ti rotor) at 35,000 x g for 30 min to pellet down the crude membrane fraction. The crude membrane pellet was then suspended in a non-denaturing lysis buffer (50 mM Tris-HCl pH 7.4, 300 mM NaCl, 5 mM EDTA and 1% TritonX-100) and processed further for co-immunoprecipitation.

### Co-immunoprecipitation

Whole tissue homogenates from forebrain and brainstem of *Tgf-β2* deficient embryos and wildtype littermates or crude membranes obtained from adult mouse whole brain were suspended in 1 mL of non-denaturing lysis buffer. Protein concentration was determined using Thermo Fisher Scientific NanoDrop 2000 spectrophotometer (at absorbance 280 nm). Immunoprecipitation was performed as described previously (Roussa et al, 2016; Pethe et al., 2023) with minor modifications. 500 µg of protein was mixed with 75 µl protein-A–Sepharose beads (Invitrogen, Carlsbad, CA; 1:1 in immunoprecipitation buffer, i.e. non-denaturing lysis buffer containing 0.1% Triton X-100) and incubated overnight at 4°C on end-to-end roller. After centrifugation for 5 min at 2300 x g, the precleared cell lysate was added to protein-A–Sepharose beads conjugated with antibody and incubated overnight at 4°C on end-to-end roller. Antibody-conjugated beads were prepared by blocking protein-A–Sepharose beads in 10% BSA in immunoprecipitation buffer and then incubating them overnight at 4°C on end-to-end roller with 5 µg of antibody (anti-KCC2 or anti-Ankyrin-G or normal mouse IgG) in immunoprecipitation buffer. The beads were centrifuged for 5 min at 2300 x g and the precleared cell lysate was added as described above. Subsequently, the beads were centrifuged, washed multiple times in immunoprecipitation buffer and the supernatant was discarded. Bound proteins and their interaction partners were resuspended in 20 µl 3× Sample buffer (62.5 mM Tris-HCl pH 6.8, 2% SDS, 10% glycine, 5% β-mercaptoethanol, 0.001% Bromophenol Blue), heated for 5 min at 95°C and immediately cooled down on ice, and processed by SDS-PAGE.

### Immunoblotting

Primary mouse hippocampal neurons at DIV4 or forebrain and brainstem tissue from E17.5 embryos were harvested and whole protein homogenates were obtained as described (Oehlke et al., 2011). Protein concentration was determined by Thermo Scientific NanoDrop 2000 spectrophotometer (absorbance at 280 nm). Electrophoresis and blotting procedures were performed as described earlier (Roussa et al., 2004). Proteins were separated by SDS-PAGE and transferred on a PVDF membrane overnight at 18V at 4°C. The following day membranes were blocked with 3% milk in TTBS (Tris buffered saline + 0.05% Tween-20) and incubated with primary antibodies overnight at 4°C. Primary antibodies were diluted as follows: anti-KCC2 1:10000, anti-GAPDH 1:20,000, polyclonal anti-Ankyrin-G 1:2000 and monoclonal anti-Ankyrin-G 1:2000 in TTBS + 0.1% milk. Subsequently, blots were incubated with horse anti-mouse or anti-rabbit IgG secondary antibodies coupled to HRP diluted 1:10,000 in TTBS + 0.1% milk and developed in enhanced chemiluminescence reagents. Signals were visualized on X-ray films. Films were scanned and protein expression was quantified by densitometrical measurement of the signal ratio Ankyrin-G: GAPDH. Differences between signal ratios for the experimental conditions were tested for statistical significance.

### Quantitative RT-PCR

Total RNA was isolated from forebrain and brainstem of *Tgf-β2^+/+^* and *Tgf-β2^-/-^* mice at E17.5 using TRIzol reagent (Invitrogen; Darmstadt, Germany) according to manufacturer’s instructions. Subsequently, 1 μg of RNA was reverse transcribed into cDNA and analyzed with quantitative real-time PCR. The following primers were used: *Ank3* primer pair 1 (Liu et al., 2017), forward (F): 5’-ACCTTGAACAGAAGCTCCTAC GCAA-3’, reverse (R): 5’-TCCGCTGCCCAACTGTAGTGTCT-3’ (GenBank accession numbers: NM_170728, NM_146005, NM_170687 - NM_170690, NM_009670, NM_170729, XM_036155521 - XM_036155557, XM_006513133, XM_006513136, XM_030244846, XM_017313778, XM_006513138, XM_006513139); *Ank3* primer pair 2 (Jenkins et al., 2015), F: 5’-GCTTTGCCTCCCTAGCTTTA-3’, R: 5’-GATATCCGTCCGCTCACAAG-3’ (GenBank accession numbers: NM_170728, NM_146005, NM_170687 - NM_170690, NM_009670, NM_170729, NM_170730, XM_036155532 - XM_036155557, XM_006513136, XM_030244846, XM_017313778, XM_006513138, XM_006513139); *Gapdh* (Ludwig et al., 2011), F: 5’- TGCACCACCAACTGCTTAGC-3’, R: 5’-TGGCATGGACTGTGGTCATG-3’ (GenBank accession number: NM_001289726). The reaction parameters were as follows: initial denaturation at 95°C for 10 min, 40 cycles of denaturation at 95°C for 15 sec, annealing at the appropriate temperature of the primer pairs for 30 sec and elongation at 72°C for 30 sec, followed by final elongation at 72°C for 10 min. All reactions were performed in triplicates and the mean Ct value was used to determine gene expression with the 2^-ΔΔCt^ method using *Gapdh* as housekeeping gene.

### Immunocytochemistry

DIV4 and DIV12 mouse primary hippocampal neurons derived from E17.5 embryos were fixed with 4% PFA for 30 min at room temperature, washed three times with PBS and permeabilized with 1% SDS for 5 min, followed by blocking with 1% BSA for 15 min. Subsequently, coverslips were incubated overnight at 4°C with primary antibodies at dilution anti-KCC2 1:250, mouse anti-Ankyrin-G 1:100, rabbit anti-Ankyrin-G 1:500 in BSA. The following day, the coverslips were washed with PBS and incubated with secondary antibodies 1:400 for 1 h. Coverslips were washed with PBS and mounted with Fluoromount-G with DAPI (4’,6’-diamidino-2-phenylindole dihydrochloride) (cat. no. 0100-20, SouthernBiotech) for nuclear staining. Double staining with primary antibodies of the same species was achieved by using the FlexAble CoraLite Plus 555 and FlexAble CoraLite 488 (cat. no. KFA002 and KFA001, Proteintech) antibody labeling kits according to manufacturer’s instructions, in order to label the antibodies with different fluorophores prior to incubation.

### Image acquisition and analysis

Images were acquired with a Leica TCS SP8 confocal microscope using a HC PL APO CS2 40x/1.30 or 63x/1.40 oil objective lens and immunofluorescence intensity was analyzed as previously described (Chleilat et al., 2020). Within each experiment, the microscope settings (laser power, detector gain, and amplifier offset) were kept the same for all scans in which the immunofluorescence intensity was compared. Z-stacks of 10– 30 optical sections with a step size of 0.4 µm were taken for at least 3 separate fields of view for each experimental condition. Maximum intensity projections were created from the z-stacks. To quantify the protein expression, LAS X software was used to select the area of interest (neuronal soma or axon initial segment; AIS) and measure the average fluorescence intensity within this area for each cell. The AIS was identified by the notable presence of Ankyrin-G immunolabeling, an extensively used marker for AIS analysis (Gutzmann et al., 2014; Leterrier et al., 2017). For measuring Ankyrin-G immunofluorescence, the AIS was traced along the axon by following the peak intensity of Ankyrin-G labeling until it was no longer visible. Background subtraction was applied to the images. Membrane localization of KCC2 and Ankyrin-G was determined by drawing a line, using the ‘Draw line’ tool of LAS X software, across the cell body and through the nucleus, spanning the periphery of the cell. The line profile tool was used to examine the fluorescence distribution across the cell. Cells expressing membrane KCC2/Ankyrin-G were defined by the presence of fluorescence intensity peaks at the periphery. Representative images for each figure were processed identically.

### Statistical analysis

Statistical analysis was performed using GraphPad Prism, Version 7.04 for Windows. Datasets were tested for normal distribution using the Shapiro-Wilk-test. Data following normal distribution were analyzed using unpaired two-tailed Student’s t-test, whereas, for datasets with non-normal distribution Mann Whitney Rank Sum Test was applied. For comparisons between more than two groups, one-way ANOVA and Tukey’s multiple comparisons test was applied. Values are shown as mean ± S.E.M. For all statistical tests, p≤0.05 was considered statistically significant.

## RESULTS

### TGF-β gain-of-function in immature hippocampal neurons has no effect on Ankyrin-G protein expression and cellular localization

Activation of TGF-β signaling in mature cortical neuronal cultures leads to an increase in Ankyrin-G protein levels and dendritic spine density (Yoon et al., 2020). However, it is not clear whether this regulation is dependent on neuronal maturation. Moreover, taken into consideration the role of isoform 2 of the TGF-β family during development, first we addressed whether TGF-β2-induced signaling regulates Ankyrin-G protein expression and cellular localization in immature hippocampal neurons. Primary neurons isolated from mouse hippocampi at E17.5, were treated at DIV4 with 2 ng/mL recombinant human TGF-β2 for 60 min and analysed by immunoblotting (Fig. 1A) and immunocytochemistry (Fig. 1B). Immunoblotting with antibody against Ankyrin-G revealed multiple immunoreactive bands corresponding to the numerous splice variants (Jenkins et al., 2013). From the known brain-specific isoforms of Ankyrin-G, only the 190 kDa one (Fig. 1A, arrow) was detected. Quantification of the ∼190 kDa band showed no significant differences after exposure of the neurons to exogenous TGF-β2 (1.27 ± 0.23 fold; n=5), compared to untreated controls (Fig. 1A). Moreover, localization of Ankyrin-G in TGF-β2-treated neurons was mainly intracellular (asterisks) and axonal (arrowheads) and was found to be similar to controls, as shown by immunofluorescence microscopy (Fig. 1B).

**Figure 1:**
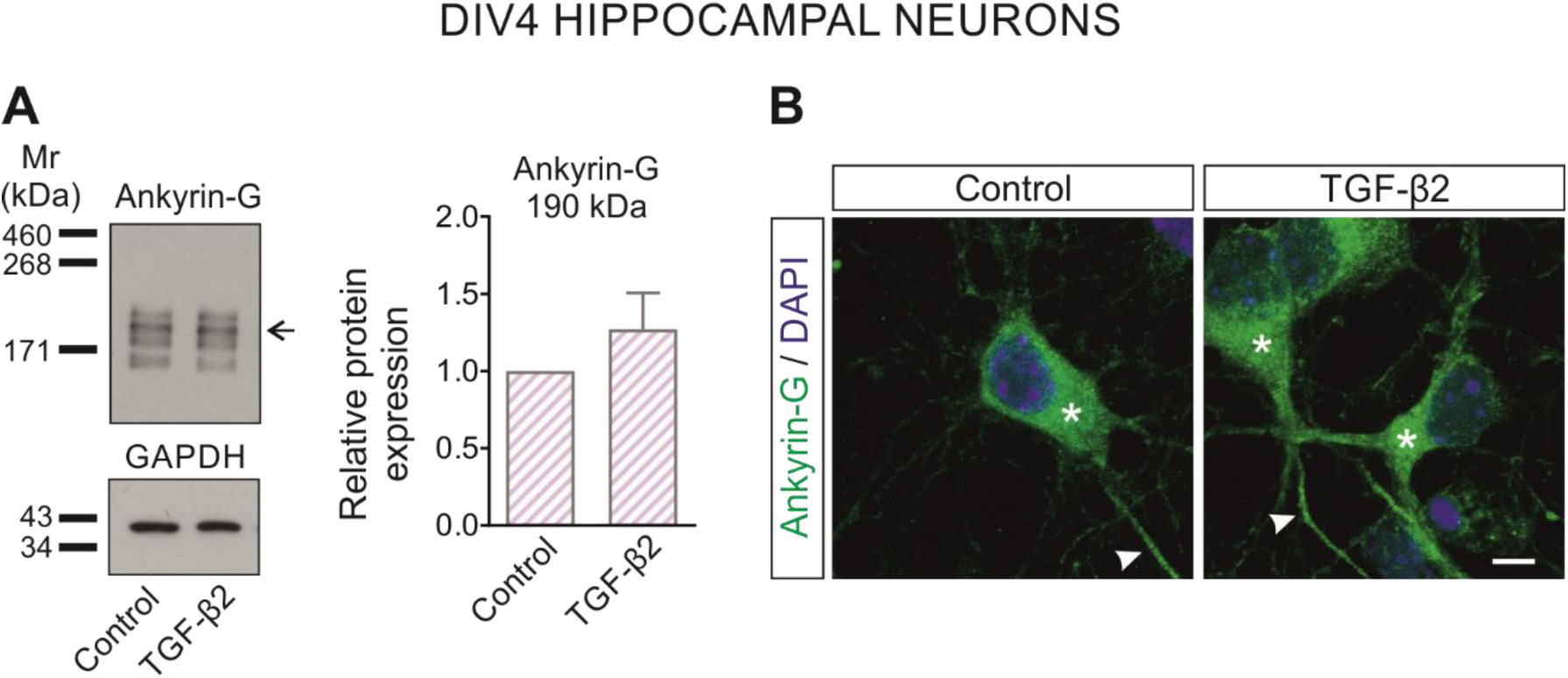
Effects of exogenous TGF-β2 on Ankyrin-G protein in mouse immature hippocampal neurons. (A) Immunoblot analysis in whole cell homogenates from immature, Day In Vitro (DIV) 4 cultured primary mouse hippocampal neurons treated with 2 ng/mL recombinant human TGF-β2 for 60 min. Arrow points to the ∼190kDa Ankyrin-G immunoreactive band. 30 µg protein was loaded per lane. Not significant, after densitometric analysis of the signal ratio Ankyrin-G 190kDa: GAPDH and two-tailed unpaired Student’s t-test, n=5. Data are shown as mean ± SEM, controls were set to 1. (B) Cellular localisation of Ankyrin-G (green) in primary immature hippocampal neuronal cultures in the presence or absence of exogenous TGF-β2 by immunofluorescence and subsequent confocal microscopy. Asterisks and arrowheads point to intracellular and axonal immunolabeling, respectively. Scale bar: 5 µm.

### Differential regulation of Ank3 transcripts in embryonic forebrain and brainstem by TGF-β2

We next investigated whether loss of the TGF-β2 ligand affects Ankyrin-G expression, using *Tgf-β2* deficient mice at E17.5. We have previously demonstrated that the impact of TGF-β2 on KCC2 varies between the embryonic forebrain and brainstem, presumably due to differences in neuronal maturation (Rigkou et al., 2022). Here, we followed the same principal to investigate a potential differential regulation of Ankyrin-G by TGF-β2 in forebrain and brainstem. We performed qRT-PCR using primer pairs that identify different transcript variants of mouse *Ankyrin-3* (*Ank3*). One primer pair recognizes all the described *Ank3* transcripts whereas qRT-PCR analysis with the second primer pair (sequences in Material and Methods) amplifies transcripts corresponding to ≤190 kDa observed molecular weight proteins.

In forebrain (Fig. 2A), expression of all *Ank3* transcript variants (1.00 ± 0.02 and 1.04 ± 0.10 for *wildtype* (*wt*; *Tgf-β2^+/+^*) and *Tgf-β2*^-/-^ respectively), as well as transcripts corresponding to ≤190 kDa (1.00 ± 0.02 and 1.04 ± 0.12 for *wt* and mutant respectively), was similar between mutants and *wt* littermates (not significant, using two-tailed unpaired Student’s *t-*test with Welch’s correction; n=6-7). In contrast, in the brainstem of *Tgf-β2^-/-^* mice (Fig. 2D), a significant downregulation in the expression of all *Ank3* transcript variants (1.00 ± 0.03 and 0.87 ± 0.07 for *wt* and mutant respectively; ^#^*p*≤0.05, using Mann-Whitney test; n=6-7), as well as of transcripts corresponding to ≤190 kDa (1.00 ± 0.01 and 0.78 ± 0.04 for *wt* and mutant respectively; ^###^*p*<0.001 using Student’s t-test; n=6-7) was detected.

**Figure 2:**
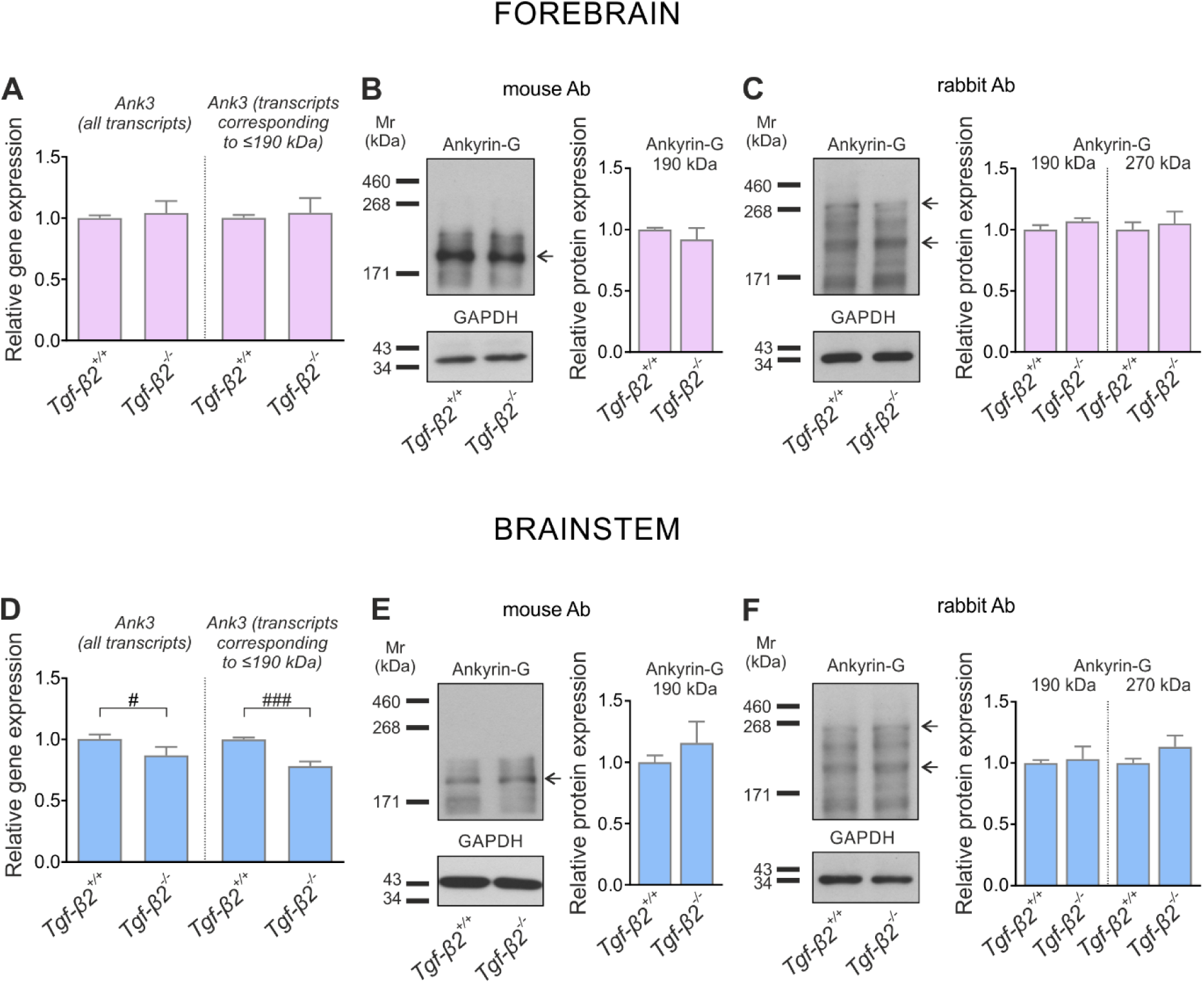
TGF-β2-dependent regulation of *Ank3* transcripts and Ankyrin-G protein in the mouse forebrain and brainstem. Quantitative RT-PCR analysis for *Ank3* transcripts on cDNA from forebrain (A) and brainstem (D) at embryonic day (E)17.5 of wildtype (*Tgf-β2^+/+^*) and *Tgf-β2^-/-^* mice (^#^*p*<0.05 and ^###^*p*<0.001 for significant decrease, using Mann-Whitney test or two-tailed unpaired Student’s *t*-test, n=6-7). Immunoblot analysis for Ankyrin-G in tissue homogenates from forebrain (B-C) and brainstem (E-F) tissue of *Tgf-β2^+/+^* and *Tgf-β2^-/-^* mice using antibody raised either in mouse (B, E) or rabbit (C, F) Quantification of immunoblots: not significant after densitometric analysis of the signal ratio Ankyrin-G: GAPDH and two-tailed unpaired Student’s *t*-test, n=3-5). The data are shown as mean ± SEM, relative to the mean of wildtype littermates. The blots are representative for three-four different litters. 30-50 µg protein was loaded per lane.

For immunoblot analysis, in order to ensure the detection of the brain-specific isoforms of Ankyrin-G, we used two different antibodies (a mouse monoclonal and a rabbit polyclonal antibody) that target different peptide sequences. Both antibodies revealed immunoreactive band at ∼190kDa (Fig. 2B, 2C, 2E, 2F, arrow). The antibody raised in rabbit however, additionally detected one of the alternative spliced giant isoforms at 270 kDa (Fig. 2C, 2F, upper arrow). In contrast, the 480 kDa isoform was not identified with either of the antibodies. Quantification of Ankyrin-G 190 kDa abundance using the mouse monoclonal antibody was comparable between *Tgf-β2* mutants and *wt* littermates in both forebrain (0.92 ± 0.09 and 1.00 ± 0.01 fold for mutant and *wt* respectively) and brainstem (1.16 ± 0.17 and 1.00 ± 0.06 fold for mutant and *wt* respectively; not significant using two-tailed unpaired Student’s *t*-test with Welch’s correction; n=6-7) (Fig. 2B, 2E). Similarly, immunoblotting using the rabbit polyclonal anti-Ankyrin-G antibody did not show any significant differences in Ankyrin-G 190 kDa protein expression either in forebrain (1.07 ± 0.03 and 1.00 ± 0.04 fold for mutant and *wt* respectively; not significant using Mann-Whitney test; n=3-6) or brainstem of *Tgf-β2^-/-^* (1.03 ± 0.10 and 1.00 ± 0.02 fold for mutant and *wt* respectively; not significant using two-tailed unpaired Student’s *t*-test; n=3-5), compared to *wt* littermates (Fig. 2C, 2F). The same results were also obtained after quantification of the 270 kDa immunoreactive band. Protein abundance of this Ankyrin-G isoform as well, showed no significant differences between *wt* and *Tgf-β2* mutants both in forebrain (Fig. 2C; 1 ± 0.06 and 1.05 ± 0.1 fold for *wt* and mutant, respectively; n=3-6) and brainstem (Fig. 2F; 1 ± 0.03 and 1.13 ± 0.09 fold for *wt* and mutant, respectively; not significant using two-tailed unpaired Student’s *t-*test; n=3-5).

These results indicate an involvement of the TGF-β2 ligand in the transcriptional expression of *Ank3* gene in embryonic brainstem. However, it remains unclear which specific variants are being regulated. Moreover, the observed transcriptional regulation does not appear to affect the protein levels of the Ankyrin-G 190 kDa and 270kDa isoforms.

### Loss of TGF-β2 impairs somatic and axonal Ankyrin-G in a neuronal maturation-dependent manner

Since total protein for the Ankyrin-G 190 kDa isoform was comparable between *wt* and *Tgf-β2* mutants, as a next step, we investigated putative TGF-β2-dependent regulation of Ankyrin-G cellular distribution in neurons at various stages of maturation. Therefore, hippocampal neurons derived from *Tgf-β2^-/-^* and *Tgf-β2^+/+^* embryos at E17.5 were cultured and subsequently analysed at either DIV4 (immature neurons) or DIV12 (differentiated neurons) using immunocytochemistry. Again, we applied the same approach as for western blot analysis (Fig. 2) and used two different anti-Ankyrin-G antibodies to potentially detect distinct labelling patterns, an assumption that turned out to be true.

As depicted in Figs 3A and 3C, confocal imaging of *Tgf-β2^+/+^* neurons at both DIV4 and DIV12, labelled with the mouse anti-Ankyrin-G antibody, revealed very prominent axonal labelling (Figs 3A, 3C; white arrowheads), confirming previous observations (Jenkins et al., 2015; Leterrier et al., 2017; Yoon et al., 2020). Moreover, intracellular Ankyrin-G labelling (Figs 3A, 3C; asterisks) was also detected. In contrast, dendritic labelling using the mouse antibody was very weak and in some cases barely detectable. Subsequently, the fluorescence intensity of Ankyrin-G within the soma and the axon of *wt* and *Tgf-β2* deficient neurons was quantified. Both at DIV4 and DIV12, somatic Ankyrin-G immunofluorescence was comparable between *wt* (Fig. 3A, 3C; 44.91 ± 0.65 and 75.83 ± 2.1 for DIV4 and DIV12 respectively) and *Tgf-β2^-/-^* (45.52 ± 0.59 and 78.56 ± 1.41 for DIV4 and DIV12 respectively; n=3-5 embryos) neurons. At DIV12, Ankyrin-G immunofluorescence within the axon was similar between *wt* and *Tgf-β2^-/-^* as well (Fig. 3C; 115.6 ± 3.03 and 109.9 ± 1.83 for *wt* and mutant respectively; not significant using two-tailed unpaired Student’s *t*-test with Welch’s correction; n=3 embryos). Interestingly, at DIV4, Ankyrin-G labelling was significantly downregulated in neurons lacking *Tgf-β2* (Fig. 3A; 33.05 ± 1.6) compared to *wt* neurons (51.81 ± 2.49; ^####^*p*<0.0001; using Mann-Whitney test; n=4 embryos).

**Figure 3:**
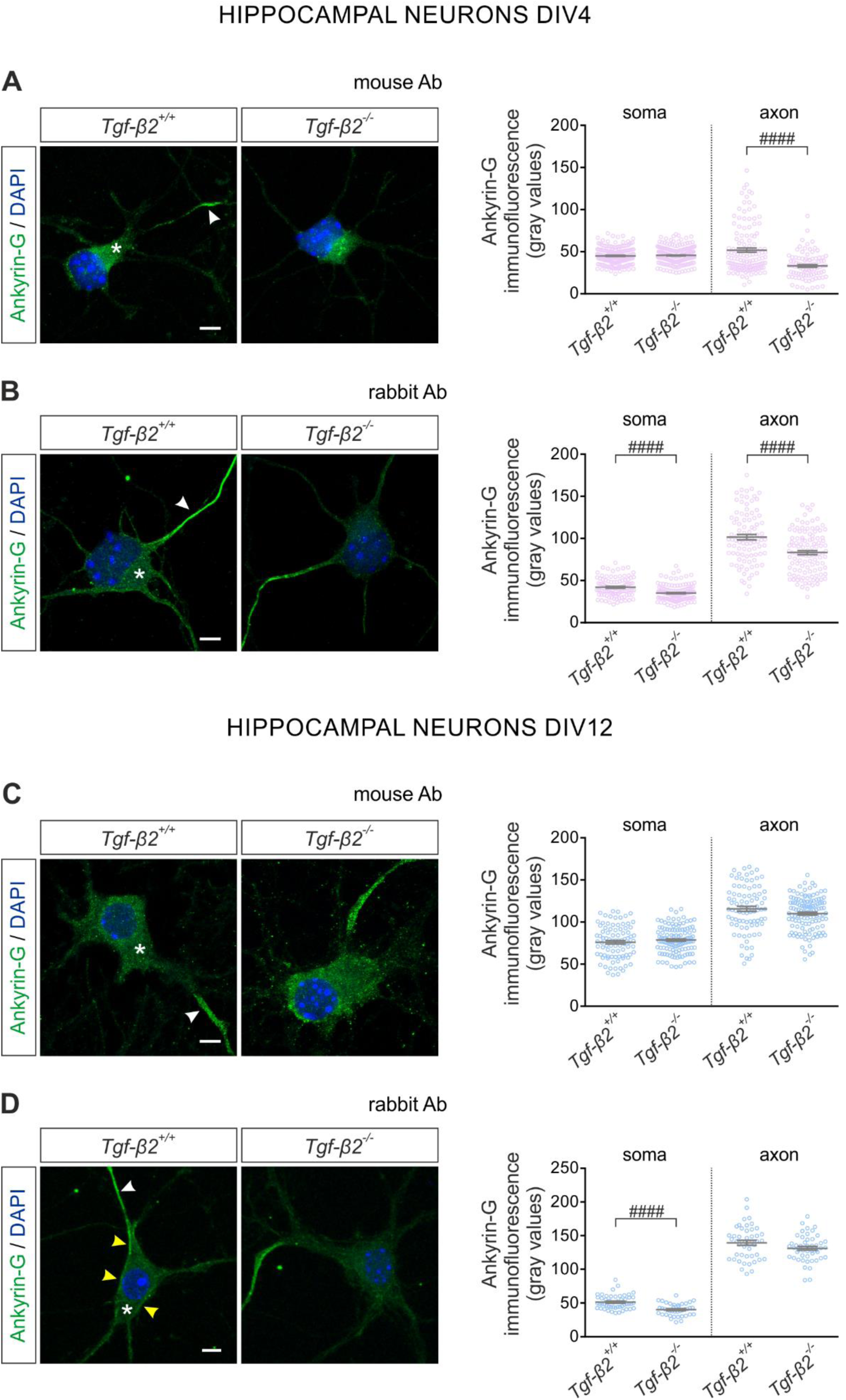
Differential TGF-β2-dependent regulation of Ankyrin-G distribution in axon and soma in immature and differentiated primary hippocampal neurons. Cellular localisation of Ankyrin-G (green) in immature (DIV4; A, B) and differentiated (DIV12; C, D) primary hippocampal neuronal cultures from *Tgf-β2^+/+^* and *Tgf-β2^-/-^* using different antibodies by immunofluorescence and subsequent confocal microscopy. Asterisks point to intracellular labelling, white arrowheads to axonal immunofluorescence and yellow arrowheads to Ankyrin-G distribution at the cell periphery, presumably plasma membrane labelling. Nuclei are labelled with DAPI. Scale bars: 5 µm. Quantification of labelling intensity: ^####^*p*<0.0001 for significant decrease in *Tgf-β2^-/-^* compared to *wildtype* littermates, using either Mann-Whitney test (A, axon, 91-128 cells from n=4 embryos; B, soma, 101-137 cells from n=3-4 embryos; D, soma, 40-50 cells from n=4-5 embryos were analysed) or two-tailed unpaired Student’s *t*-test with Welch’s correction (B, axon, 91-111 from 3-4 embryos; C, soma/ axon, 86-123 cells from n=3-4 embryos were analysed). Data are presented as mean ± SEM.

When employing the rabbit anti-Ankyrin-G antibody, prominent labelling across the axon (Figs 3B, 3D; white arrowheads) and additionally, dendritic and plasma membrane labelling was detected, as indicated by yellow arrowheads (Figs 3B, 3D). Quantification of labelling intensity at DIV4 obtained with the rabbit anti-Ankyrin-G antibody, showed a significant downregulation within the soma and the axon of *Tgf-β2^-/-^* neurons (Fig. 3B; 35.11 ± 0.77 and 83.18 ± 2.41 for soma and axon, respectively), compared to neurons from *wt* littermates (42.02 ± 1.09 and 101.5 ± 3.25 for soma and axon, respectively; ^####^*p*<0.0001 using Mann-Whitney test or two-tailed unpaired Student’s *t*-test with Welch’s correction; n=3-4 embryos). In differentiating neurons at DIV12, however, while axonal Ankyrin-G intensity was comparable between wt and *Tgf-β2^-/-^* neurons (Fig. 3D; 139.4 ± 3.67 and 131.2 ± 2.83 for *wt* and mutant respectively; not significant using two-tailed unpaired Student’s *t*-test), Ankyrin-G immunofluorescence intensity within the soma was significantly downregulated in *Tgf-β2^-/-^* compared to *wt* neurons (Fig. 3D; 51.35 ± 1.47 and 40.1 ± 1.43 for *wt* and mutant respectively; ^####^*p*<0.0001 using Mann-Whitney test; n=4-5 embryos). Moreover, as depicted in Fig. 3D, *wt* neurons displayed notable membrane Ankyrin-G labeling (highlighted by yellow arrowheads) which was absent in *Tgf-β2* deficient neurons, implicating that the downregulation in somatic Ankyrin-G immunofluorescence might be partly attributed to impaired membrane abundance of Ankyrin-G.

Collectively, these data demonstrate distinct Ankyrin-G labelling patterns in neurons stained with different antibodies, probably reflecting detection of different Ankyrin-G isoforms. Moreover, the results suggest that TGF-β2 may orchestrate axon-specific expression and somatic membrane abundance of Ankyrin-G in immature and differentiating neurons, respectively, *in vitro*.

### Evidence for a Golgi-associated Ankyrin-G isoform in neurons

As shown in Fig. 3, with both anti-Ankyrin-G antibodies used, intracellular Ankyrin-G immunolabelling was detectable. Notably, using the mouse monoclonal antibody, two distinct labelling patterns could be identified: neurons revealing diffuse intracellular Ankyrin-G labelling (Figs 3A, 3C, asterisk) and neurons exhibiting a Golgi-associated Ankyrin-G labelling pattern (Fig. 4, arrows) were present within the same culture. This observation is particularly interesting, since previous studies have identified an alternative transcript of *Ank3* in the kidney, namely AnkG119, as an isoform that binds and co-localises with β1 spectrin in the ER and Golgi apparatus (Devarajan et al., 1996). To verify Ankyrin-G localization in the Golgi, double immunofluorescence for Ankyrin-G and the Golgi marker GolgA5 was performed. The results are presented in Fig. 4. At DIV4 (Fig. 4A), in both *Tgf-β2^+/+^* and *Tgf-β2* mutants, GolgA5 immunofluorescence revealed a predominant cup-like pattern that co-localised with Ankyrin-G immunofluorescence. At DIV12 (Fig. 4B), a more diffuse perinuclear labelling pattern could be observed for both GolgA5 and Ankyrin-G and showed no differences between genotypes. Notably, using the rabbit antibody, Golgi-associated Ankyrin-G labelling was undetectable.

**Figure 4:**
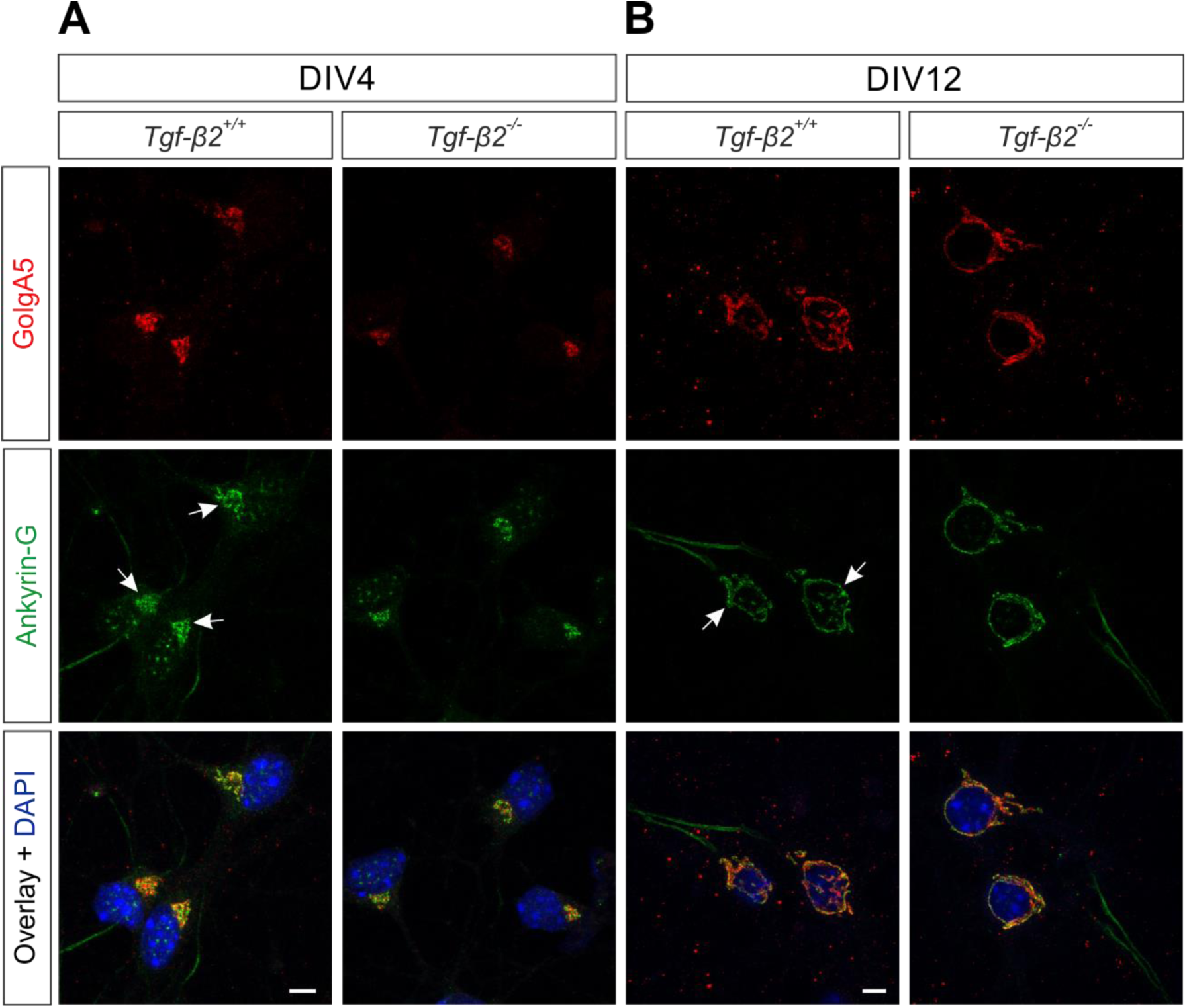
Golgi-associated Ankyrin-G isoform is not regulated by TGF-β2. Double immunofluorescence for the Golgi marker GolgA5 (red) and Ankyrin-G (green) in immature (DIV4, A) and differentiated (DIV12, B) primary hippocampal neurons in *wildtype* (*Tgf-β2*^+/+^) and *Tgf-β2* deficient mice, followed by confocal microscopy. Co-localisation of the proteins was observed that was comparable between the genotypes. Arrows point to Golgi-associated Ankyrin-G labelling. Nuclei are labelled with DAPI. Scale bar: 5 µm.

These observations provide a hint for a putative Golgi-associated Ankyrin-G isoform in neurons.

### KCC2 and Ankyrin-G are TGF-β2-dependent interaction partners in the embryonic brainstem

In a native interactome study from adult mouse forebrain plasma membranes, Ankyrin-G was reported to be in the top 150 proteins associated with KCC2 immunocomplex (Smalley et al., 2020b). We sought to assess whether KCC2 and Ankyrin-G interact *in vivo* in adult mouse brains. Therefore, crude membranes isolated from whole adult mouse brains were subjected to co-immunoprecipitation using beads coated with antibodies against KCC2 or Ankyrin-G followed by immunoblotting, while beads coated with normal mouse IgG were used as a negative control. Beads coated with anti-KCC2 or anti-Ankyrin-G antibody were successfully able to immunoprecipitate and enrich their respective proteins (Fig. 5A, Fig. 5B, lane 3, arrows), compared to the input fraction (Fig. 5A, 5B, lane 1). Normal mouse IgG neither pulled down KCC2 (Fig. 5A, lane 2) or Ankyrin-G (Fig. 5B, lane 2). Anti-KCC2 beads were also successful in pulling down the 190 kDa isoform of Ankyrin-G protein (Fig. 5C, lane 3, arrow), while the IgG fraction remained clean (Fig. 5C, lane 2). Similarly, co-immunoprecipitation by beads coated with anti-Ankyrin-G antibodies successfully precipitated KCC2 as well (Fig. 5D, lane 3, arrow), with the IgG fraction being clean (Fig. 5D, lane 2). These data show that KCC2 and Ankyrin-G are indeed part of the same immunocomplex, in the adult mouse brain.

**Figure 5:**
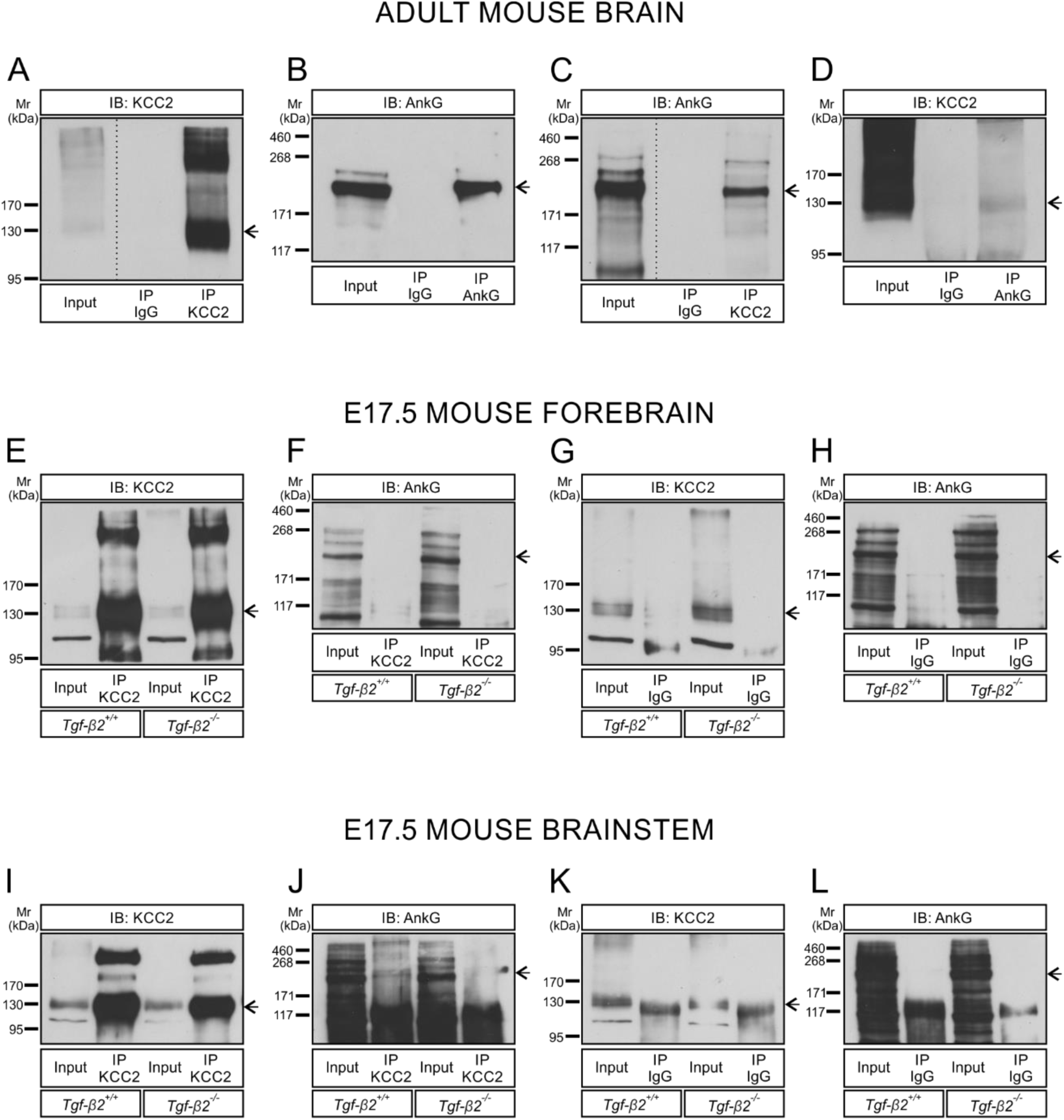
Ankyrin-G and KCC2 are interaction partners *in vivo* and regulated by TGF-β2 in embryonic brainstem. (A-D) Co-immunoprecipitation of KCC2 and Ankyrin-G (AnkG) in crude membranes from whole mouse brain. Enrichment of KCC2 (A) and AnkG (B) following immunoprecipitation (IP) of the respective proteins. Antibodies against KCC2 (C) and AnkG (D), but not IgG (C, D) were able to immunoprecipitate AnkG and KCC2, respectively, as detected by immunoblotting. (E-H) Immunoprecipitation of KCC2 from whole tissue homogenates of embryonic day 17.5 *Tgf-β2^+/+^* and *Tgf-β2^-/-^* mouse forebrain. Enrichment of KCC2 (E) following IP with KCC2 but not with IP IgG (G). Antibody against KCC2 (F) was not able to immunoprecipitate AnkG. (I-L) Co-immunoprecipitation of KCC2 and AnkG in homogenates from brainstem of embryonic day 17.5 *Tgf-β2^+/+^* and *Tgf-β2^-/-^* mice. KCC2 enrichment following IP with anti-KCC2 antibody (I), but not with IgG (K). Antibodies against KCC2 (J) were able to precipitate AnkG from wildtype but not *Tgf-β2^-/-^* brainstem lysate, while IgG (L), did not immunoprecipitate AnkG. Arrows point to the relevant protein bands; n=2 for A-H; n=5 for I-L. Dotted line indicates where the same blot has been cut to eliminate empty lanes. 500µg protein (input) or IP derived from 500µg protein was loaded.

Having established Ankyrin-G/KCC2 interaction in adult mouse brains, next, the question was addressed whether KCC2/Ankyrin-G interaction takes place in the embryonic brain as well and if this interaction might be regulated by TGF-β2. Therefore, co-immunoprecipitation experiments were performed in whole protein derived from forebrain or brainstem tissue from E17.5 *wildtype* and *Tgf-β2* mutants. First, we confirmed that the pull-down assay was successful as KCC2 was enriched in the immunoprecipitated fraction (Fig. 5E, lanes 2 and 4, arrow; Fig. 5I, lanes 2 and 4, arrow) of all experimental conditions, compared to the respective inputs (Fig. 5E, lanes 1 and 3; Fig. 5I, lanes 1 and 3, arrow). The negative controls using IgG, on the other hand, did not show the KCC2 specific band (Fig. 5G, lanes 2 and 4, arrow; Fig, 5K, lanes 2 and 4, arrow). Although an immunoreactive band was obtained in the IgG fractions from the brainstem (Fig. 5K, lanes 2 and 4; Fig. 5L, lanes 2 and 4), this band was considered non-specific as it was obtained following immunoblotting with both antibodies against KCC2 (Fig. 5K) or against Ankyrin-G (Fig. 5L).

Interestingly, in forebrain, as shown in Fig. 5F, IP with KCC2 antibody did not immunoprecipitate Ankyrin-G (Fig. 5F, arrow), neither in *wildtype* (Fig. 5F, lane 2) nor in *Tgf-β2^-/-^* (Fig. 5F, lane 4). However, in the brainstem, co-immunoprecipitation with anti-KCC2 antibodies successfully pulled down the 190 kDa Ankyrin-G, as indicated by the arrow in Fig. 5J, in *wildtype* embryos (Fig. 5J, lane 2), but not in *Tgf-β2* null mutants (Fig. 5J, lane 4).

These data provide a strong hint for TGF-β2-mediated interaction of KCC2 with Ankyrin-G, and particularly with the 190 kDa isoform, in the differentiating neurons of the brainstem.

### Rescue of impaired membrane Ankyrin-G in Tgf-β2 deficient neurons by exogenous TGF-β2

We next asked whether KCC2 co-localizes with Ankyrin-G either in dendrites or other subcellular compartments in DIV12 neuronal cultures, and if yes, whether this co-localization is regulated by TGF-β2. Previous studies have established that KCC2 resides at the somatodendritic plasma membrane of differentiating and mature neurons (Williams et al., 1999; Gulyás et al., 2001). With this in mind, differentiating neurons at DIV12 were co-stained with antibody against KCC2 and the rabbit polyclonal anti-Ankyrin-G antibody, shown to recognize plasma membrane and dendritic Ankyrin-G (Fig. 3D). Fig. 6A shows that in *Tgf-β2^+/+^* neurons both KCC2 (a1) and Ankyrin-G (a1’) were indeed present at the somatic plasma membrane (white arrowheads), and prominent membrane co-localization was evident (yellow arrowheads, a1’’). In contrast, *Tgf-β2*-deficient neurons exhibited a notable absence of KCC2/Ankyrin-G membrane co-localization due to either reduced or even undetectable KCC2 (b1) and Ankyrin-G (b1’), compared to *wt* cells (n=4-5 embryos). Importantly, lack of KCC2/Ankyrin-G membrane co-localization in TGF-β2 deficient neurons was rescued (yellow arrowheads in b2’’) in the presence of exogenous TGF-β2, thus highlighting the TGF-β2-dependency of membrane targeting of both proteins (26 cells from n=2 embryos were analysed). Exposure of *wt* neurons to exogenous TGF-β2 had no effect on Ankyrin-G immunolocalization (Fig. 6A, a2’, white arrowheads) but led to an increase in KCC2 membrane abundance (Fig. 6A, a2; white arrowheads), as previously reported (Roussa et al., 2016). These results are further illustrated by representative line scans (Fig. 5C) from *Tgf-β2^+/+^ and Tgf-β2^-/-^* neurons in the presence or absence of exogenous TGF-β2. Peaks of KCC2 (red) and Ankyrin-G (green) immunofluorescence (arrows) were not detected in neurons of *Tgf-β2* deficient animals.

**Figure 6:**
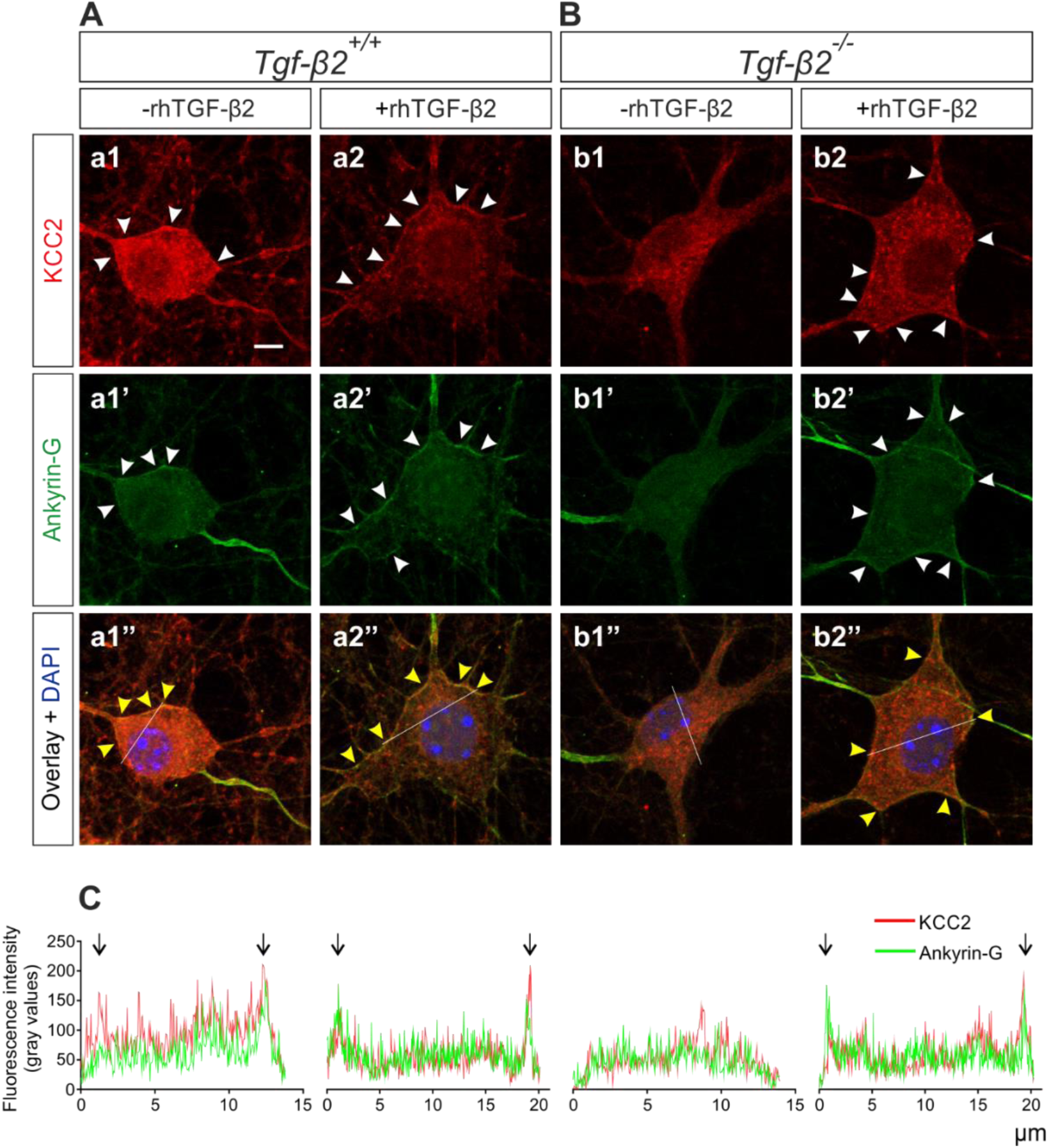
Rescue of impaired membrane Ankyrin-G in *Tgf-β2* deficient neurons by exogenous TGF-β2. Double immunofluorescence for KCC2 (red; a1-b2) and Ankyrin-G (green; a1’-b2’) in cultured differentiated hippocampal neurons from wildtype (*Tgf-β2^+/+^*; a1, a2) and *Tgf-β2* deficient (*Tgf-β2^-/-^*; b1, b2) in the presence (2ng/ml; a2, b2) or absence (a1, b1) of exogenous recombinant human (rh) TGF-β2 for 60 min. Nuclei are stained with DAPI. White arrowheads point to plasma membrane labeling, yellow arrowheads indicate increased or rescued plasma membrane labeling in the presence of exogenous TGF-β2. Scale bar: 5 µm. (C): line scans for KCC2 (red) and Ankyrin-G (green) immunofluorescence. Arrows point to peaks of fluorescence intensity at the periphery of the neurons.

These results suggest that putative interaction of KCC2 with Ankyrin-G at the neuronal plasma membrane of differentiating neurons is regulated by TGF-β2.

### Knock-down of Ankyrin-G impairs membrane KCC2

To further explore putative interdependency between the two interaction partners, either *Kcc2b* (Fig. 7D, 7I) or *Ank3* (Fig. 7E, 7J) were transiently knocked down in cultured hippocampal neurons at DIV11 using specific siRNA (as described in Material and Methods) and subsequently processed for immunocytochemistry for KCC2 (Fig. 7A-7E) and Ankyrin-G (Fig. 7F-7J), followed by quantification of the fluorescence intensity within the soma (intracellular and plasma membrane) (Fig 7K, 7L). Cells transfected with negative control siRNA (Fig. 7C, 7H), cells treated only with the transfection reagent (mock; Fig. 7B, 7G) and non-transfected cells (Fig. 7A, 7F) were used as controls. Green arrows point to cells successfully transfected with the different siRNAs, whereas non-transfected cells from the same coverslip are pointed by white arrows. Quantification of KCC2 fluorescence intensity showed no differences between cells transfected with negative control siRNA (Fig. 7C, 7K; 60.26 ± 3.20, n=23 cells) and cells treated only with either the transfection reagent [mock (Fig. 7B, 7K; 59.83 ± 3.63; n=19 cells)] or culture medium [non-transfected (Fig. 7A, 7K; 60.46 ± 1.79; n=55 cells)]. Similarly, Fig 7L shows that Ankyrin-G labeling intensity was comparable between non-transfected (Fig. 7F; 74.84 ± 2.13; n=64 cells), mock (Fig. 7G; 76.91 ± 2.86; n=25 cells) and negative siRNA transfected (Fig. 7H; 67.61 ± 3.48; n=26 cells) neurons. On the contrary, KCC2 and Ankyrin-G immunofluorescence were significantly downregulated in cells transfected with siRNA against *Kcc2* (Fig. 7D; 46.65 ± 2.13, n=12 cells; ^##^*p*<0.01) and siRNA against *Ank3* (Fig. 7J; 52.18 ± 1.70. n=48 cells; ^####^*p*<0.0001), respectively, compared to their respective non-transfected neurons (using one-way ANOVA with Tukey’s multiple comparisons test), thereby confirming the specificity of the targeted siRNAs and the efficiency of the knockdown on KCC2 and Ankyrin-G protein levels. Interestingly, while transfection with si*Kcc2* had no effect on Ankyrin-G labeling (Fig. 7I, 7L; 70.98 ± 2.17, n=11 cells), somatic KCC2 fluorescence intensity was significantly downregulated after transfection with siRNA against *Ank3* (Fig. 7E, 7K; 44.21 ± 1.58, n=33 cells; ^####^*p*<0.0001 using one-way ANOVA with Tukey’s multiple comparisons test), compared to non-transfected cells. As a next step, the question was addressed whether *Ank3* knockdown regulates the intracellular or plasma membrane fraction of KCC2. Fig. 7M highlights a cell transfected with si*Ank3* and an adjacent non-transfected cell labeled for KCC2.

**Figure 7:**
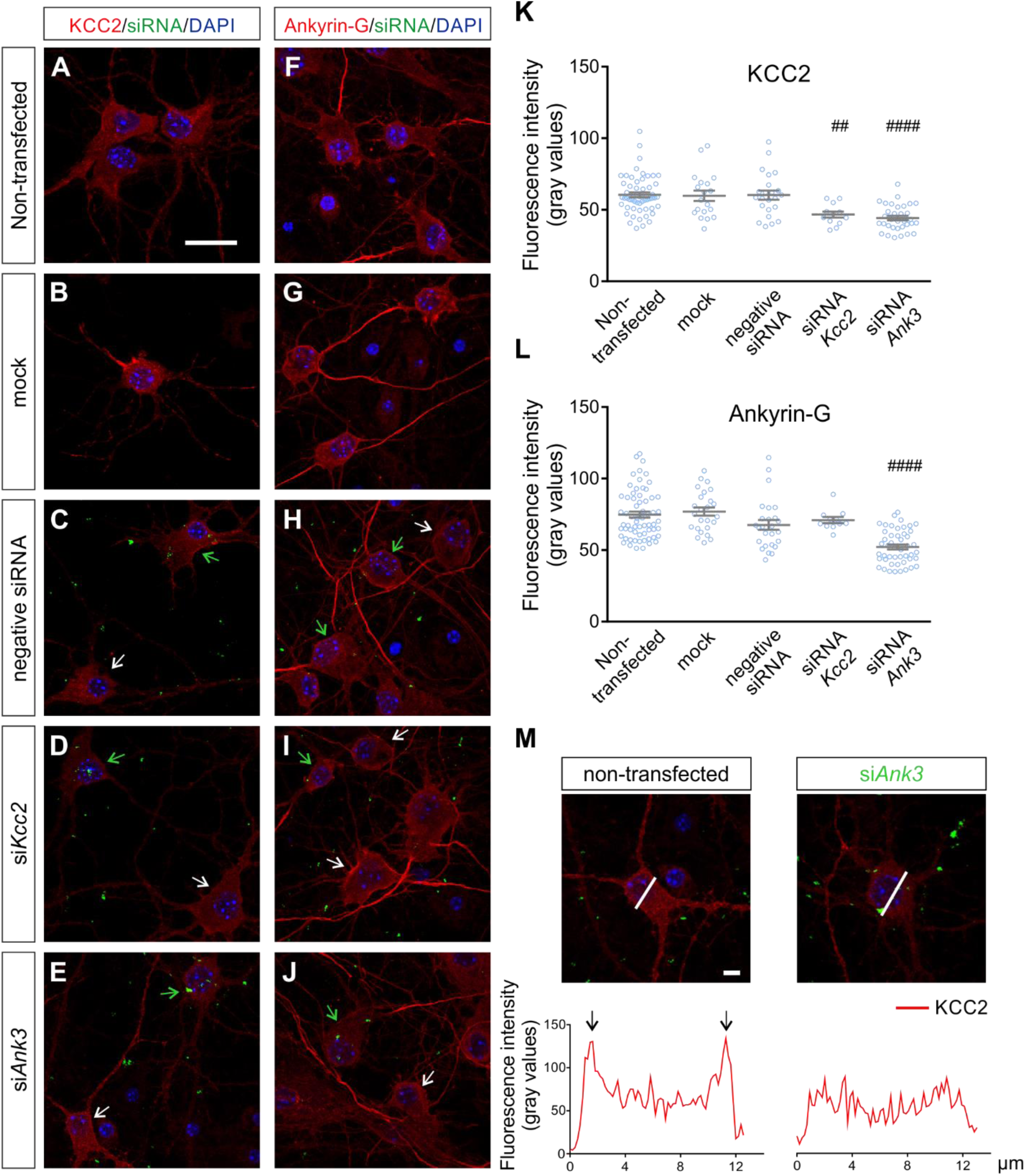
Regulation of membrane KCC2 by Ankyrin-G. Immunofluorescence for KCC2 (red, A-E) and Ankyrin-G (red, F-J) in differentiated primary hippocampal neurons cultured in normal culture medium (non-transfected; A, F), or in transfection reagent (mock; B, G), or transfected with negative siRNA (green; C, H), specific siRNA against *Kcc2* (si*Kcc2*; green; D, I) or specific siRNA against *Ank3* (si*Ank3*; green; E, J) and subsequent confocal microscopy. Nuclei are labeled with DAPI. White arrows point to non-transfected neurons, green arrows indicate cells successfully transfected with the respective siRNA. (K, L): Quantification of fluorescence intensity (^##^*p*<0.01 and ^####^*p*<0.0001 for significant decrease, compared to non-transfected cells, using one-way ANOVA with Tukey’s multiple comparisons test; 11-55 cells were analysed per experimental condition. Values are shown as mean ± SEM. Scale bar: 20 µm. (M): representative line scans following immunofluorescence for KCC2 (red) in a non-transfected neuron (left) and a neuron transfected with si*Ank3* (green; right). Arrows point to spikes of fluorescence intensity at the periphery of the cell in the non-transfected neuron. A total of ∼25 cells were analysed for each experimental condition. Nuclei were labelled with DAPI (blue). Scale bar: 5 µm.

Somatic KCC2 immunofluorescence was clearly reduced in the neuron transfected with si*Ank3,* compared to non-transfected cell. Moreover, representative line scan of the non-transfected neuron shows intensity peaks of KCC2 immunofluorescence at the cell periphery (Fig. 7M, arrows) presumably the area of the plasma membrane, whereas in line scan of the si*Ank3* transfected cell intensity peaks were not documented.

Taken together, these data suggest a potential role of Ankyrin-G in maintaining KCC2 protein levels or in translocating KCC2 to the membrane or both.

## DISCUSSION

KCC2 plays a crucial role in neuronal development and function by regulating chloride homeostasis and dendritic spine formation. Disruption of any of these processes can lead to brain-related pathologies, therefore, it is imperative to understand the mechanisms driving KCC2 expression and function in order to identify putative therapeutic targets.

We have previously introduced TGF-β2 as a key regulator of KCC2 throughout neuronal maturation and identified mechanisms through which TGF-β2 exerts its effect on KCC2: In immature neurons, TGF-β2 regulates *Kcc2* expression via binding of AP2β to *Kcc2* promoter, whereas in differentiated neurons TGF-β2 regulates phosphorylation of KCC2 at T1007 (Rigkou et al., 2022), as well as KCC2 trafficking to the membrane via Rab11b (Roussa et al., 2016). Here, we provide evidence for a putative novel mechanism of TGF-β2 -dependent regulation of KCC2, by regulating its interaction with the scaffolding protein Ankyrin-G.

We formulated the hypothesis that TGF-β2 regulates KCC2/Ankyrin-G interaction. This hypothesis was based on the following observations: first, TGF-β2 regulates KCC2 at the transcriptional and post-translational level, as mentioned above, second, TGF-β signaling regulates Ankyrin-G via Usp9X (Yoon et al., 2020), and third, a recent interactome has shown Ankyrin-G as KCC2 interaction partner (Smalley et al., 2020b).

### Ankyrin-G regulation by TGF-β2 signaling

To test this hypothesis, as a first step, a potential role of TGF-β2 signaling in Ankyrin-G regulation was investigated. Gain-of-function and loss-of-function experiments were employed by inducing signaling upon treatment of hippocampal neurons with exogenous TGF-β2 and by using neuronal cultures and tissue from *Tgf-β2* constitutive knockout mice, respectively. *Ank3* gene codes for several Ankyrin-G isoforms, most of which are poorly characterized (reviewed by Yoon et al., 2022). Hence, we focused on the isoforms with well-established roles in neurons, known as Ankyrin-G 190/ 270/ 480 kDa (Smith et al., 2014; Jenkins et al., 2015). Treating immature hippocampal neurons with exogenous TGF-β2 did not affect the expression of Ankyrin-G 190 kDa protein (Fig. 1). This lack of effect can be attributed to the presence of endogenous TGF-β2, which may saturate the signaling pathway, similar to what was previously observed on KCC2 expression (Rigkou et al., 2022). Subsequent loss-of-function experiments in brain areas at different maturation stages in *Tgf-β2* mutants and expression analysis of multiple transcript variants of *Ank3*, showed no regulation in the forebrain of *Tgf-β2* mutants (Fig. 2A), consistent with the immunoblotting results (Figs 2B, 2C). However, these observations do not exclude a potential regulation of other isoforms, such as Ankyrin-G 480 kDa, which could not be detected by immunoblotting. In contrast, in the brainstem, where, at E17.5, neurons are considered more mature than those in the forebrain, a reduction in *Ank3* transcript expression in *Tgf-β2* mutants was observed. Surprisingly, Ankyrin-G 190 kDa and 270 kDa protein abundance remained unchanged (Figs 2E, 2F), suggesting that other isoform(s) may be regulated by TGF-β2. In the present study, two different commercially available antibodies against Ankyrin-G were used that likely recognize distinct Ankyrin-G isoforms. Several lines of evidence support this view: First, distinct pattern of immunoreactive bands in western blot could be obtained, i.e. the rabbit polyclonal anti-Ankyrin-G antibody revealed immunoreactive bands at 190 and 270 kDa Ankyrin-G, the mouse monoclonal antibody recognized only a 190 kDa band. Moreover, whereas both antibodies showed prominent axonal and intracellular labeling, plasma membrane labeling was only evident when the rabbit anti-Ankyrin-G was used (Fig. 3D), and Golgi-associated Ankyrin-G was exclusively detected with the mouse monoclonal anti-Ankyrin-G antibody. (Fig. 4). Smaller Ankyrin-G isoforms that lack the membrane-binding domain were found in endosomes and lysosomes of fibroblasts (Ignatiuk et al., 2006), whereas Golgi-associated Ankyrin-G of 119 kDa was identified in kidney and muscle (Devarajan et al., 1996), however, as of our knowledge, the presence of a Golgi-associated Ankyrin-G isoform in neurons has not been previously documented. At this point it should be noted that Golgi-associated Ankyrin-G labelling was consistent but not exclusive using the mouse monoclonal anti-Ankyrin-G antibody. Indeed, the labeling pattern of Ankyrin-G, even when employing the same antibody, exhibited variability between different cultures or even within the same culture. Since primary hippocampal cultures contain a mixture of neurons at various developmental stages that may not be synchronized, this observation suggests the possibility of differential expression patterns for multiple Ankyrin-G isoforms during neuronal development. The existence of numerous *Ank3* transcript variants increases the complexity of studying the particular protein and underlines the necessity for the development of additional tools such as isoform-specific antibodies, to enable more precise investigation of these protein variants. One significant finding of the present work is the downregulation of axonal Ankyrin-G immunofluorescence in immature (DIV4) *Tgf-β2* deficient neurons (Fig. 3A, 3B). This implicates that TGF-β2 may play a role in axon organization, aligning with previous research proposing that TGF-β signaling is essential for axon specification during early mouse brain development (Yi et al., 2010). However, the observation that TGF-β2 regulates axonal Ankyrin-G at DIV4 but not at DIV12 implicates putative compensatory mechanisms by other TGF-β isoforms. An alternative explanation would imply that several Ankyrin-G isoforms are expressed in the axon and/or the isoform expressed at DIV4 is developmentally downregulated at DIV12. Indeed Ankyrin-G isoforms are known to be differentially regulated after birth: axonal Ankyrin-G 480 is downregulated and dendritic Ankyrin-G 190 is upregulated, whereas the Ankyrin-G 270 remains at a constant level (Yoshimura et al., 2016).

In differentiated neurons, quantification of somatic Ankyrin-G immunofluorescence showed a significant downregulation in *Tgf-β2* deficient neurons labeled with the rabbit but not the mouse anti-Ankyrin-G antibody (Fig. 3D). This discrepancy can be simply explained by the detection of membrane-bound Ankyrin-G only when employing the rabbit antibody. Importantly, the decrease in somatic Ankyrin-G intensity in cells lacking *Tgf-β2*, can be attributed to the absence of membrane-localized Ankyrin-G, thus indicating a pivotal role of TGF-β2 in regulating Ankyrin-G membrane abundance in differentiating neurons.

In summary, TGF-β2 modulates *Ank3* expression in the brainstem and influences axonal and membrane localization of Ankyrin-G in immature and differentiating neurons, respectively. The specific transcript variants undergoing regulation require further investigation. Our results highlight the multifaceted regulation of Ankyrin-G by TGF-β2 during neuronal development.

### TGF-β2-mediated interaction of KCC2 with Ankyrin-G

The rapid functional changes of KCC2 under physiological conditions and in response to pathological stimuli, primarily result from post-translational events rather than transcriptional regulation of the transporter (Puskarjov et al., 2012). KCC2 membrane trafficking, stability and functionality is critically influenced by protein-protein interactions. Consequently, an increasing number of studies focus on characterizing KCC2 interactome (Mahadevan et al., 2017; Smalley et al., 2020a and 2020b; Awabdh et al., 2022). Many interaction partners of KCC2 have been identified, some of them have been thoroughly investigated. Transport function of KCC2 is promoted by its interaction with brain-type creatine kinase (CKA) (Inoue et al., 2004; 2006), Ras-associated binding protein 11b (Rab11b) (Roussa et al., 2016) and Gephyrin (Awabdh et al., 2022), while PACSIN1 is proposed as a negative regulator of KCC2 expression and function (Mahadevan et al., 2017). Interaction of KCC2 with the kainate receptor subunit 2 (GluK2) enhances its membrane expression and function (Pressey et al., 2017) and additionally contributes in the structural maturation of dendritic spines (Kesaf, Khirug et al., 2020). The non-canonical function of KCC2 is mediated through interactions with the cytoskeleton-associated protein 4.1N (Li et al., 2007), which also interacts with GluK2 (Copits and Swanson, 2013), and the guanine nucleotide exchange factor β-PIX (Llano et al., 2015).

It was demonstrated that KCC2 interacts with Ankyrin-G in adult mouse forebrain plasma membrane preparations (Smalley et al., 2020b). However, the specific isoform(s) of Ankyrin-G involved in this interaction remain unclear. As discussed above, axon-specific Ankyrin-G is regulated by TGF-β2, however, the absence of KCC2 in the axon (Szabadics et al., 2006) turns axon-associated isoforms less likely candidates for interaction. The results demonstrate that TGF-β2 regulates membrane Ankyrin-G (Figs 3 and 6), which could be a promising candidate interaction partner since KCC2 is typically found in the somatodendritic plasma membrane of mature neurons. Although total protein of Ankyrin-G 190 kDa isoform, which is enriched in dendrites (Smith et al., 2014), does not appear to be regulated by TGF-β2 (Fig. 2), this does not preclude the possibility of a TGF-β2-dependent interaction with KCC2. Support for a TGF-β2-dependent KCC2/Ankyrin-G interaction was obtained by co-immunoprecipitation experiments performed in forebrain and brainstem of *wildtype* and *Tgf-β2^-/-^* mice at E17.5 (Fig. 5). While in embryonic forebrains, no interaction between KCC2 and Ankyrin-G in either *wildtype* or *Tgf-β2* deficient mice could be detected, in the brainstem, KCC2 interacts with Ankyrin-G 190 kDa in *wildtype* mice, but this interaction is absent in *Tgf-β2* mutants. Considering that KCC2 interacts with Ankyrin-G in both the adult forebrain (Smalley et al., 2020b) and embryonic brainstem, where KCC2 is present in the neuronal membrane, and not in the embryonic forebrain, where membrane-bound KCC2 is not yet expressed, it is reasonable to assume that this interaction primarily occurs in the neuronal membrane and/or dendrites. Along this line, double immunofluorescence for KCC2 and Ankyrin-G revealed co-localization in the somatic plasma membrane in differentiated hippocampal neurons of *wildtype* but not in *Tgf-β2* deficient neurons, an effect that could be rescued by exogenous TGF-β2 (Fig. 6). Since the phenotype in *Tgf-β2* deficient neurons could not be compensated by endogenous expression of other TGF-β isoforms, i.e. TGF-β3, these results point to an isoform-specific effect of TGF-β2 on Ankyrin-G. However, caution should be maintained in interpreting the results, since it is known that TGF-β2 regulates trafficking and membrane expression of KCC2 as well (Roussa et al., 2016), and therefore an impaired KCC2/Ankyrin-G co-localization at the membrane of the mutant neurons would be expected even without any effect of TGF-β2 on membrane Ankyrin-G abundance. The novel and relevant finding here is the rescue of membrane Ankyrin-G following exposure of the deficient neurons to exogenous TGF-β2. Of course, co-localization alone does not definitively prove a TGF-β2-dependent interaction between KCC2 and Ankyrin-G in the plasma membrane. Although tempting, the conclusive link between TGF-β2 and KCC2/Ankyrin-G interaction at the neuronal membrane is still missing. Additional investigations, are therefore required to provide conclusive evidence.

To shed light into the significance of the interaction between KCC2 and Ankyrin-G and understand putative interdependency of these two proteins, we conducted RNAi experiments targeting either *Kcc2* or *Ank3*. While Ankyrin-G protein was not affected when *Kcc2* was transiently knocked down, *Ank3* knockdown resulted in a notable decrease in KCC2 protein abundance, especially in membrane-bound KCC2. This reduction in the membrane pool of KCC2 could be an outcome of total protein downregulation or impaired surface stability, however, again further experiments are needed necessary to elucidate whether the intracellular or surface pool of KCC2 is affected by *Ank3* depletion. In both scenarios, we speculate that Ankyrin-G may operate either as a direct downstream effector of TGF-β2 or indirectly through a TGF-β-induced interaction with Usp9X (Yoon et al., 2020). This putative mechanism might serve to prevent KCC2 from proteasomal degradation and promote its stability at the cell surface and would represent an additional TGF-β2-mediated regulatory mechanism on membrane KCC2.

In conclusion, the present work provides new insights into Ankyrin-G regulation by TGF-β2 during neuronal maturation and first evidence of a TGF-β2-mediated interaction of KCC2 with Ankyrin-G.

## REFERENCES

Awabdh, S. Al, Donneger, F., Goutierre, M., Séveno, M., Vigy, O., Weinzettl, P., Russeau, M., Moutkine, I., Lévi, S., Marin, P., Poncer, J. C. (2022). Gephyrin Interacts with the K-Cl Cotransporter KCC2 to Regulate Its Surface Expression and Function in Cortical Neurons. The Journal of Neuroscience, 42(2), 166. 10.1523/JNEUROSCI.2926-20.2021

Bi, C., Wu, J., Jiang, T., Liu, Q., Cai, W., Yu, P., Cai, T., Zhao, M., Jiang, Y., Sun, Z. S. (2012). Mutations of ANK3 identified by exome sequencing are associated with autism susceptibility. Human Mutation, 33(12), 1635–1638. 10.1002/humu.22174

Chleilat, E., Pethe, A., Pfeifer, D., Krieglstein, K., Roussa, E. (2020). TGF-β Signaling Regulates SLC8A3 Expression and Prevents Oxidative Stress in Developing Midbrain Dopaminergic and Dorsal Raphe Serotonergic Neurons. International Journal of Molecular Sciences, 21(8), 2735. 10.3390/ijms21082735

Copits, B. A., Swanson, G. T. (2013). Kainate Receptor Post-translational Modifications Differentially Regulate Association with 4.1N to Control Activity-dependent Receptor Endocytosis*. Journal of Biological Chemistry, 288(13), 8952–8965. 10.1074/jbc.M112.440719

Devarajan, P., Stabach, P. R., Mann, A. S., Ardito, T., Kashgarian, M., Morrow, J. S. (1996). Identification of a small cytoplasmic ankyrin (AnkG119) in the kidney and muscle that binds beta I sigma spectrin and associates with the Golgi apparatus. Journal of Cell Biology, 133(4), 819–830. 10.1083/jcb.133.4.819

Dünker, N., Krieglstein, K. (2000). Targeted mutations of transforming growth factor-β genes reveal important roles in mouse development and adult homeostasis. European Journal of Biochemistry, 267(24), 6982–6988. 10.1046/j.1432-1327.2000.01825.x

Ferreira, M. A. R., O’Donovan, M. C., Meng, Y. A., Jones, I. R., Ruderfer, D. M., Jones, L., Fan, J., Kirov, G., Perlis, R. H., Green, E. K., Smoller, J. W., Grozeva, D., Stone, J., Nikolov, I., Chambert, K., Hamshere, M. L., Nimgaonkar, V. L., Moskvina, V., Thase, M. E., et al. (2008). Collaborative genome-wide association analysis supports a role for ANK3 and CACNA1C in bipolar disorder. Nature Genetics, 40(9), 1056–1058. 10.1038/ng.209

Fiumelli, H., Briner, A., Puskarjov, M., Blaesse, P., Belem, B. J. T., Dayer, A. G., Kaila, K., Martin, J.-L., Vutskits, L. (2013). An Ion Transport-Independent Role for the Cation-Chloride Cotransporter KCC2 in Dendritic Spinogenesis In Vivo. Cerebral Cortex, 23(2), 378–388. 10.1093/cercor/bhs027

Gulyás, A. I., Sík, A., Payne, J. A., Kaila, K., Freund, T. F. (2001). The KCl cotransporter, KCC2, is highly expressed in the vicinity of excitatory synapses in the rat hippocampus. European Journal of Neuroscience, 13(12), 2205–2217. 10.1046/j.0953-816x.2001.01600.x

Gutzmann, A., Erguel, N., Grossmann, R., Schultz, C., Wahle, P., Engelhardt, M. (2014). A period of structural plasticity at the axon initial segment in developing visual cortex. Frontiers in Neuroanatomy, 8. https://www.frontiersin.org/articles/10.3389/fnana.2014.00011

Heupel, K., Sargsyan, V., Plomp, J. J., Rickmann, M., Varoqueaux, F., Zhang, W., Krieglstein, K. (2008). Loss of transforming growth factor-beta 2 leads to impairment of central synapse function. Neural Development, 3(1), 25. 10.1186/1749-8104-3-25

Ignatiuk, A., Quickfall, J. P., Hawrysh, A. D., Chamberlain, M. D., Anderson, D. H. (2006). The Smaller Isoforms of Ankyrin 3 Bind to the p85 Subunit of Phosphatidylinositol 3’-Kinase and Enhance Platelet-derived Growth Factor Receptor Down-regulation *. Journal of Biological Chemistry, 281(9), 5956–5964. 10.1074/jbc.M510032200

Inoue, K., Ueno, S., Fukuda, A. (2004). Interaction of neuron-specific K+-Cl− cotransporter, KCC2, with brain-type creatine kinase. FEBS Letters, 564(1–2), 131–135. 10.1016/S0014-5793(04)00328-X

Inoue, K., Yamada, J., Ueno, S., Fukuda, A. (2006). Brain-type creatine kinase activates neuron-specific K+-Cl– co-transporter KCC2. Journal of Neurochemistry, 96(2), 598–608. 10.1111/j.1471-4159.2005.03560.x

Jaenisch, N., Witte, O. W., Frahm, C. (2010). Downregulation of Potassium Chloride Cotransporter KCC2 After Transient Focal Cerebral Ischemia. Stroke, 41(3), e151–e159. 10.1161/STROKEAHA.109.570424

Jenkins, P. M., Kim, N., Jones, S. L., Tseng, W. C., Svitkina, T. M., Yin, H. H., Bennett, V. (2015). Giant ankyrin-G: A critical innovation in vertebrate evolution of fast and integrated neuronal signaling. Proceedings of the National Academy of Sciences, 112(4), 957–964. 10.1073/pnas.1416544112

Jenkins, P. M., Vasavda, C., Hostettler, J., Davis, J. Q., Abdi, K., Bennett, V. (2013). E-cadherin Polarity Is Determined by a Multifunction Motif Mediating Lateral Membrane Retention through Ankyrin-G and Apical-lateral Transcytosis through Clathrin. Journal of Biological Chemistry, 288(20), 14018–14031. 10.1074/jbc.M113.454439

Kahle, K. T., Merner, N. D., Friedel, P., Silayeva, L., Liang, B., Khanna, A., Shang, Y., Lachance-Touchette, P., Bourassa, C., Levert, A., Dion, P. A., Walcott, B., Spiegelman, D., Dionne-Laporte, A., Hodgkinson, A., Awadalla, P., Nikbakht, H., Majewski, J., Cossette, P., Deeb, T. Z., Moss, S. J., Medina, I., Rouleau, G. A. (2014). Genetically encoded impairment of neuronal KCC2 cotransporter function in human idiopathic generalized epilepsy. EMBO Reports, 15(7), 766–774. 10.15252/embr.201438840

Kesaf, S., Khirug, S., Dinh, E., Saez Garcia, M., Soni, S., Orav, E., Delpire, E., Taira, T., Lauri, S. E., Rivera, C. (2020). The Kainate Receptor Subunit GluK2 Interacts With KCC2 to Promote Maturation of Dendritic Spines. Frontiers in Cellular Neuroscience, 14. https://www.frontiersin.org/articles/10.3389/fncel.2020.00252

Krieglstein, K., Zheng, F., Unsicker, K., Alzheimer, C. (2011). More than being protective: functional roles for TGF-β/activin signaling pathways at central synapses. Trends in Neurosciences, 34(8), 421–429. 10.1016/j.tins.2011.06.002

Lacmann, A., Hess, D., Gohla, G., Roussa, E., Krieglstein, K. (2007). Activity-dependent release of transforming growth factor-beta in a neuronal network *in vitro*. Neuroscience, 150(3), 647–657. 10.1016/j.neuroscience.2007.09.046

Leterrier, C., Clerc, N., Rueda-Boroni, F., Montersino, A., Dargent, B., Castets, F. (2017). Ankyrin G Membrane Partners Drive the Establishment and Maintenance of the Axon Initial Segment. Frontiers in Cellular Neuroscience, 11. https://www.frontiersin.org/articles/10.3389/fncel.2017.00006

Li, H., Khirug, S., Cai, C., Ludwig, A., Blaesse, P., Kolikova, J., Afzalov, R., Coleman, S. K., Lauri, S., Airaksinen, M. S., Keinänen, K., Khiroug, L., Saarma, M., Kaila, K., Rivera, C. (2007). KCC2 Interacts with the Dendritic Cytoskeleton to Promote Spine Development. Neuron, 56(6), 1019–1033. 10.1016/j.neuron.2007.10.039

Liu, C., Zhang, L., Wu, J., Sui, X., Xu, Y., Huang, L., Han, Y., Zhu, H., Li, Y., Sun, X., Qin, C. (2017). AnkG hemizygous mice present cognitive impairment and elevated anxiety/depressive-like traits associated with decreased expression of GABA receptors and postsynaptic density protein. Experimental Brain Research, 235(11), 3375–3390. 10.1007/s00221-017-5056-7

Llano, O., Smirnov, S., Soni, S., Golubtsov, A., Guillemin, I., Hotulainen, P., Medina, I., Nothwang, H. G., Rivera, C., Ludwig, A. (2015). KCC2 regulates actin dynamics in dendritic spines via interaction with β-PIX. Journal of Cell Biology, 209(5), 671–686. 10.1083/jcb.201411008

Ludwig, A., Uvarov, P., Soni, S., Thomas-Crusells, J., Airaksinen, M. S., Rivera, C. (2011). Early Growth Response 4 Mediates BDNF Induction of Potassium Chloride Cotransporter 2 Transcription. The Journal of Neuroscience, 31(2), 644. 10.1523/JNEUROSCI.2006-10.2011

Mahadevan, V., Khademullah, C. S., Dargaei, Z., Chevrier, J., Uvarov, P., Kwan, J., Bagshaw, R. D., Pawson, T., Emili, A., De Koninck, Y., Anggono, V., Airaksinen, M., Woodin, M. A. (2017). Native KCC2 interactome reveals PACSIN1 as a critical regulator of synaptic inhibition. ELife, 6, e28270. 10.7554/eLife.28270

Mapplebeck, J. C. S., Lorenzo, L.-E., Lee, K. Y., Gauthier, C., Muley, M. M., De Koninck, Y., Prescott, S. A., Salter, M. W. (2019). Chloride Dysregulation through Downregulation of KCC2 Mediates Neuropathic Pain in Both Sexes. Cell Reports, 28(3), 590–596.e4. 10.1016/j.celrep.2019.06.059

Mavrovic, M., Uvarov, P., Delpire, E., Vutskits, L., Kaila, K., Puskarjov, M. (2020). Loss of non-canonical KCC2 functions promotes developmental apoptosis of cortical projection neurons. EMBO Reports, 21(4), e48880. 10.15252/embr.201948880

Merner, N. D., Chandler, M. R., Bourassa, C., Liang, B., Khanna, A. R., Dion, P., Rouleau, G. A., Kahle, K. T. (2015). Regulatory domain or CpG site variation in SLC12A5, encoding the chloride transporter KCC2, in human autism and schizophrenia. Frontiers in Cellular Neuroscience, 9. https://www.frontiersin.org/articles/10.3389/fncel.2015.00386

Meyers, E. A., Kessler, J. A. (2017). TGF-β Family Signaling in Neural and Neuronal Differentiation, Development, and Function. Cold Spring Harbor Perspectives in Biology, 9(8), a022244. 10.1101/cshperspect.a022244

Oehlke, O., Martin, H. W., Osterberg, N., Roussa, E. (2011). Rab11b and its effector Rip11 regulate the acidosis-induced traffic of V-ATPase in salivary ducts. Journal of Cellular Physiology, 226(3), 638–651. 10.1002/jcp.22388

Pethe, A., Hamze, M., Giannaki, M., Heimrich, B., Medina, I., Hartmann, A.-M., Roussa, E. (2023). K^+^/Cl^−^ cotransporter 2 (KCC2) and Na^+^/HCO3^−^ cotransporter 1 (NBCe1) interaction modulates profile of KCC2 phosphorylation. Frontiers in Cellular Neuroscience, 17. https://www.frontiersin.org/articles/10.3389/fncel.2023.1253424

Pressey, J. C., Mahadevan, V., Khademullah, C. S., Dargaei, Z., Chevrier, J., Ye, W., Huang, M., Chauhan, A. K., Meas, S. J., Uvarov, P., Airaksinen, M. S., Woodin, M. A. (2017). A kainate receptor subunit promotes the recycling of the neuron-specific K^+^-Cl^-^ co-transporter KCC2 in hippocampal neurons. Journal of Biological Chemistry, 292(15), 6190–6201. 10.1074/jbc.M116.767236

Puskarjov, M., Ahmad, F., Kaila, K., Blaesse, P. (2012). Activity-Dependent Cleavage of the K-Cl Cotransporter KCC2 Mediated by Calcium-Activated Protease Calpain. The Journal of Neuroscience, 32(33), 11356. 10.1523/JNEUROSCI.6265-11.2012

Puskarjov, M., Seja, P., Heron, S. E., Williams, T. C., Ahmad, F., Iona, X., Oliver, K. L., Grinton, B. E., Vutskits, L., Scheffer, I. E., Petrou, S., Blaesse, P., Dibbens, L. M., Berkovic, S. F., Kaila, K. (2014). A variant of KCC2 from patients with febrile seizures impairs neuronal Cl− extrusion and dendritic spine formation. EMBO Reports, 15(6), 723– 729. 10.1002/embr.201438749

Rigkou, A., Magyar, A., Speer, J. M., & Roussa, E. (2022). TGF-β2 Regulates Transcription of the K^+^/Cl^−^ Cotransporter 2 (KCC2) in Immature Neurons and Its Phosphorylation at T1007 in Differentiated Neurons. Cells, 11(23), 3861. 10.3390/cells11233861

Rivera, C., Voipio, J., Payne, J. A., Ruusuvuori, E., Lahtinen, H., Lamsa, K., Pirvola, U., Saarma, M., Kaila, K. (1999). The K+/Cl− co-transporter KCC2 renders GABA hyperpolarizing during neuronal maturation. Nature, 397(6716), 251–255. 10.1038/16697

Roussa, E., Nastainczyk, W., Thévenod, F. (2004). Differential expression of electrogenic NBC1 (SLC4A4) variants in rat kidney and pancreas. Biochemical and Biophysical Research Communications, 314(2), 382–389. 10.1016/j.bbrc.2003.12.099

Roussa, E., Speer, J. M., Chudotvorova, I., Khakipoor, S., Smirnov, S., Rivera, C., Krieglstein, K. (2016). The membrane trafficking and functionality of the K^+^-Cl^−^ co-transporter KCC2 is regulated by TGF-β2. Journal of Cell Science, 129(18), 3485–3498. 10.1242/jcs.189860

Saitsu, H., Watanabe, M., Akita, T., Ohba, C., Sugai, K., Ong, W. P., Shiraishi, H., Yuasa, S., Matsumoto, H., Beng, K. T., Saitoh, S., Miyatake, S., Nakashima, M., Miyake, N., Kato, M., Fukuda, A., Matsumoto, N. (2016). Impaired neuronal KCC2 function by biallelic SLC12A5 mutations in migrating focal seizures and severe developmental delay. Scientific Reports, 6(1), 30072. 10.1038/srep30072

Sanford, L. P., Ormsby, I., Groot, A. C. G., Sariola, H., Friedman, R., Boivin, G. P., Cardell, E. Lou, Doetschman, T. (1997). TGFβ2 knockout mice have multiple developmental defects that are non-overlapping with other TGFβ knockout phenotypes. Development, 124(13), 2659–2670. 10.1242/dev.124.13.2659

Schulze, T. G., Detera-Wadleigh, S. D., Akula, N., Gupta, A., Kassem, L., Steele, J., Pearl, J., Strohmaier, J., Breuer, R., Schwarz, M., Propping, P., Nöthen, M. M., Cichon, S., Schumacher, J., Rietschel, M., McMahon, F. J., NIMH Genetics Initiative Bipolar Disorder Consortium, Rietschel, M., McMahon, F. J. (2009). Two variants in Ankyrin 3 (ANK3) are independent genetic risk factors for bipolar disorder. Molecular Psychiatry, 14(5), 487–491. 10.1038/mp.2008.134

Smalley, J. L., Kontou, G., Choi, C., Ren, Q., Albrecht, D., Abiraman, K., Santos, M. A. R., Bope, C. E., Deeb, T. Z., Davies, P. A., Brandon, N. J., Moss, S. J. (2020a). Isolation and Characterization of Multi-Protein Complexes Enriched in the K-Cl Co-transporter 2 From Brain Plasma Membranes. Frontiers in Molecular Neuroscience, 13. https://www.frontiersin.org/articles/10.3389/fnmol.2020.563091

Smalley, J. L., Kontou, G., Choi, C., Ren, Q., Albrecht, D., Abiraman, K., Santos, M. A. R., Bope, C. E., Deeb, T. Z., Davies, P. A., Brandon, N. J., Moss, S. J. (2020b). The K-Cl co-transporter 2 is a point of convergence for multiple autism spectrum disorder and epilepsy risk gene products. BioRxiv, 2020.03.02.973859. 10.1101/2020.03.02.973859

Smith, K. R., Kopeikina, K. J., Fawcett-Patel, J. M., Leaderbrand, K., Gao, R., Schürmann, B., Myczek, K., Radulovic, J., Swanson, G. T., Penzes, P. (2014). Psychiatric Risk Factor ANK3/Ankyrin-G Nanodomains Regulate the Structure and Function of Glutamatergic Synapses. Neuron, 84(2), 399–415. 10.1016/j.neuron.2014.10.010

Stödberg, T., McTague, A., Ruiz, A. J., Hirata, H., Zhen, J., Long, P., Farabella, I., Meyer, E., Kawahara, A., Vassallo, G., Stivaros, S. M., Bjursell, M. K., Stranneheim, H., Tigerschiöld, S., Persson, B., Bangash, I., Das, K., Hughes, D., Lesko, N., et al. (2015). Mutations in SLC12A5 in epilepsy of infancy with migrating focal seizures. Nature Communications, 6(1), 8038. 10.1038/ncomms9038

Szabadics, J., Varga, C., Molnár, G., Oláh, S., Barzó, P., Tamás, G. (2006). Excitatory Effect of GABAergic Axo-Axonic Cells in Cortical Microcircuits. Science, 311(5758), 233–235. 10.1126/science.1121325

Tyzio, R., Nardou, R., Ferrari, D. C., Tsintsadze, T., Shahrokhi, A., Eftekhari, S., Khalilov, I., Tsintsadze, V., Brouchoud, C., Chazal, G., Lemonnier, E., Lozovaya, N., Burnashev, N., Ben-Ari, Y. (2014). Oxytocin-Mediated GABA Inhibition During Delivery Attenuates Autism Pathogenesis in Rodent Offspring. Science, 343(6171), 675–679. 10.1126/science.1247190

Unsicker, K., Flanders, K. C., Cissel, D. S., Lafyatis, R., Sporn, M. B. (1991). Transforming growth factor beta isoforms in the adult rat central and peripheral nervous system. Neuroscience, 44(3), 613–625. 10.1016/0306-4522(91)90082-Y

Williams, J. R., Sharp, J. W., Kumari, V. G., Wilson, M., Payne, J. A. (1999). The Neuron-specific K-Cl Cotransporter, KCC2: ANTIBODY DEVELOPMENT AND INITIAL CHARACTERIZATION OF THE PROTEIN*. Journal of Biological Chemistry, 274(18), 12656–12664. 10.1074/jbc.274.18.12656

Yi, J. J., Barnes, A. P., Hand, R., Polleux, F., Ehlers, M. D. (2010). TGF-β Signaling Specifies Axons during Brain Development. Cell, 142(1), 144–157. 10.1016/j.cell.2010.06.010

Yoon, S., Parnell, E., Penzes, P. (2020). TGF-β-Induced Phosphorylation of Usp9X Stabilizes Ankyrin-G and Regulates Dendritic Spine Development and Maintenance. Cell Reports, 31(8), 107685. 10.1016/j.celrep.2020.107685

Yoon, S., Piguel, N. H., Penzes, P. (2022). Roles and mechanisms of ankyrin-G in neuropsychiatric disorders. Experimental & Molecular Medicine, 54(7), 867–877. 10.1038/s12276-022-00798-w

Yoshimura, T., Stevens, S. R., Leterrier, C., Stankewich, M. C., Rasband, M. N. (2016). Developmental changes in expression of βIV spectrin splice variants at axon initial segments and nodes of Ranvier. Frontiers in Cellular Neuroscience. 10, 304. doi: 10.3389/fncel.2016.00304

